# Targeting *Ogt* in ADPKD mitigates metabolic reprogramming and renal cystogenesis, extending survival

**DOI:** 10.64898/2025.12.14.693989

**Authors:** Matthew A. Kavanaugh, Saleem Ahmad, Dona Greta Isai, Heather A.L. Riddell, Amanda P. Assis, Chadhve Ranganathan, Aakriti Chaturvedi, Vincent Lam, Nikhitha Muthineni, Rayyan Abid, Jenna E. Jurgensmeyer, Casey Blades, Michele T. Pritchard, Madhulika Sharma, Darren P. Wallace, Stephen C. Parnell, Chad Slawson, Pamela V. Tran

**Affiliations:** Dept. of Cell Biology and Physiology, The Jared Grantham Kidney Institute, University of Kansas Medical Center, Kansas City, KS; Dept. of Biochemistry and Molecular Biology, The Jared Grantham Kidney Institute, University of Kansas Medical Center, Kansas City, KS; Dept of Cancer Biology, University of Kansas Medical Center, Kansas City, KS; Dept. of Pharmacology, Toxicology and Therapeutics, The Jared Grantham Kidney Institute, University of Kansas Medical Center, Kansas City, KS; Dept. of Internal Medicine, The Jared Grantham Kidney Institute, University of Kansas Medical Center, Kansas City, KS

## Abstract

Aberrant cell metabolism drives autosomal dominant polycystic kidney disease (ADPKD). O-GlcNAcylation, a metabolically regulated post-translational modification, is elevated in ADPKD kidneys. Using rapidly and slowly progressive ADPKD mouse models, we demonstrate that deleting O-GlcNAc transferase (*Ogt*) reduces renal cystogenesis and extends survival in a rapidly progressive model from postnatal day 21 to over a year. Pharmacological OGT inhibition similarly reduced cyst formation of patient-derived renal epithelial cells *in vitro*. In *Pkd1* conditional knockout kidneys, *Ogt* deletion maintained phosphorylated AMPK and mitochondrial respiratory chain complex levels, preserving cellular energy sensing and production. Further, metabolomic analysis revealed normalization of glycolysis and of the hexosamine and hyaluronic acid biosynthesis pathways. In contrast, dysregulation of these pathways in *Pkd1* conditional knockout kidneys culminated in increased tricarboxylic acid cycle entry, increased O-GlcNAc, and increased hyaluronic acid in the extracellular matrix, respectively. These findings identify *Ogt* as a central metabolic regulator and therapeutic target, linking metabolism to intracellular and extracellular mechanisms of cyst formation.

## Introduction

Autosomal Dominant Polycystic Kidney Disease (ADPKD) affects approximately 1/1000 individuals worldwide and is the 4^th^ leading cause of renal failure worldwide. ADPKD is caused by mutation of *PKD1* or *PKD2*, which encode polycystin 1 (PC1) and 2 (PC2) proteins, respectively. PC1 and PC2 form a heteromeric complex that is thought to function as a receptor-ion channel complex in various cellular compartments, including the primary cilium. Mutation of either *PKD1* or *PKD2* results in increased renal epithelial proliferation and fluid secretion, driving formation of fluid-filled cysts. Cysts can initiate as early as *in utero* and grow progressively over multiple decades. Cysts enlarge kidney size and compress surrounding parenchyma, causing injury and fibrosis, which lead to kidney failure typically in the 6^th^ decade of life.

Over the last decade, altered cellular metabolism has emerged as an important driver of ADPKD progression^1–4^. Altered glucose metabolism^1,5–7^, increased glutamine consumption^4,8^, reduced activation of AMP-activated protein kinase (AMPK)^3,9^, and decreased mitochondrial function^10,11^ are some of the metabolic alterations that characterize the disease. Notably, correction of these metabolic alterations in preclinical models attenuates the disease^1–4,9–11^.

A pathway that may integrate these metabolic processes is the nutrient-sensitive Hexosamine Biosynthetic Pathway (HBP). The HBP is initiated by glucose and utilizes ATP, glutamine, acetyl-Coenzyme A, and uridine to produce uridine-5’-diphosphate-N-Acetylglucosamine (UDP-GlcNAc)^12,13^. UDP-GlcNAc is transferred onto serine and threonine residues of protein substrates as O-linked N-Acetylglucosamine (O-GlcNAc). Addition of O-GlcNAc onto a protein target can regulate the subcellular localization and activity of the protein. Two enzymes, O-GlcNAc transferase (OGT) and O-GlcNAcase (OGA), add and remove the O-GlcNAc moiety, respectively^12,13^. Thus, the HBP ties energy metabolism to protein post-translational modification (PTM).

Oscillations in any of the HBP nutrient inputs (glucose, ATP, glutamine, acetyl-CoA, uridine) will alter pathway flux^12,13^. Chronic metabolic stress, caused by nutrient deprivation or excess, increases HBP activity and O-GlcNAcylation. In metabolic diseases, such as cardiovascular disease, diabetes, and cancer, chronic activation of the HBP increasing O-GlcNAcylation is detrimental^14–20^. Increased O-GlcNAcylation causes enhanced glycolysis, reduced mitochondrial oxidative phosphorylation, and reduced AMPK activity^21–27^. We have shown that O-GlcNAc and OGT are increased in cyst-lining renal epithelial cells of juvenile and adult mouse ADPKD kidney tissue and that global levels of O-GlcNAc are increased in mouse ADPKD kidney lysates^28^. Here we investigate a functional role for increased O-GlcNAcylation in mouse ADPKD kidneys and extend our analyses to human ADPKD tissue and primary renal epithelial cells. We further examine the metabolic effects of *Ogt* deletion in mouse ADPKD kidneys and identify *Ogt* as a key metabolic regulator controlling both intracellular and extracellular processes of cyst formation.

## Results

### *Ogt* deletion in a rapidly progressive ADPKD mouse model attenuates disease severity and increases survival

To determine whether increased O-GlcNAcylation is pathogenic in a rapidly progressive ADPKD mouse model, we deleted *Pkd1* and/or *Ogt* using the *HoxB7-Cre* recombinase. *HoxB7* is expressed in the embryonic mesonephric duct and ureteric bud, which differentiates into the collecting duct and ureter. At postnatal day (P)14 and P21, deletion of *Ogt* alone resulted in kidneys that appeared largely normal, but with some dilations that did not affect kidney function (**Fig 1A-1F; Fig S1**). Deletion of *Pkd1* caused renal cystogenesis, increased kidney weight:body weight (KW/BW) ratios, and increased blood urea nitrogen (BUN) levels, indicating impaired kidney function. Relative to *Pkd1* cko mice, *Pkd1;Ogt* double knockout (dko) mice showed markedly reduced KW/BW ratios, renal cystogenesis and BUN. Moreover, in contrast to *Pkd1* cko mice which died between P13-P21, one out of nine *Pkd1;Ogt* double knock-out (dko) mice survived to P110 and remaining dko mice (8/9) survived beyond one year of age along with control and *Ogt* cko littermates **(Fig 1G)**. Three *Pkd1;Ogt* dko mice showed hydronephrosis at 1 year of age, suggesting a potential combined role for *Pkd1* and *Ogt* in the ureter. However, 5 *Pkd1;Ogt* dko mice at 1 year of age did not develop hydronephrosis and had KW/BW ratios lower than those of dko mice at P14 (**Fig 1H**). Additionally, BUN levels of these 5 *Pkd1;Ogt* dko mice at one year of age were similar to those of dko mice at P14 (**Fig 1I**). This indicates that after P14, the drastically attenuating effects of *Ogt* deletion on ADPKD enabled body growth to outpace kidney growth, and kidney function to remain stable.

**Figure 1.**
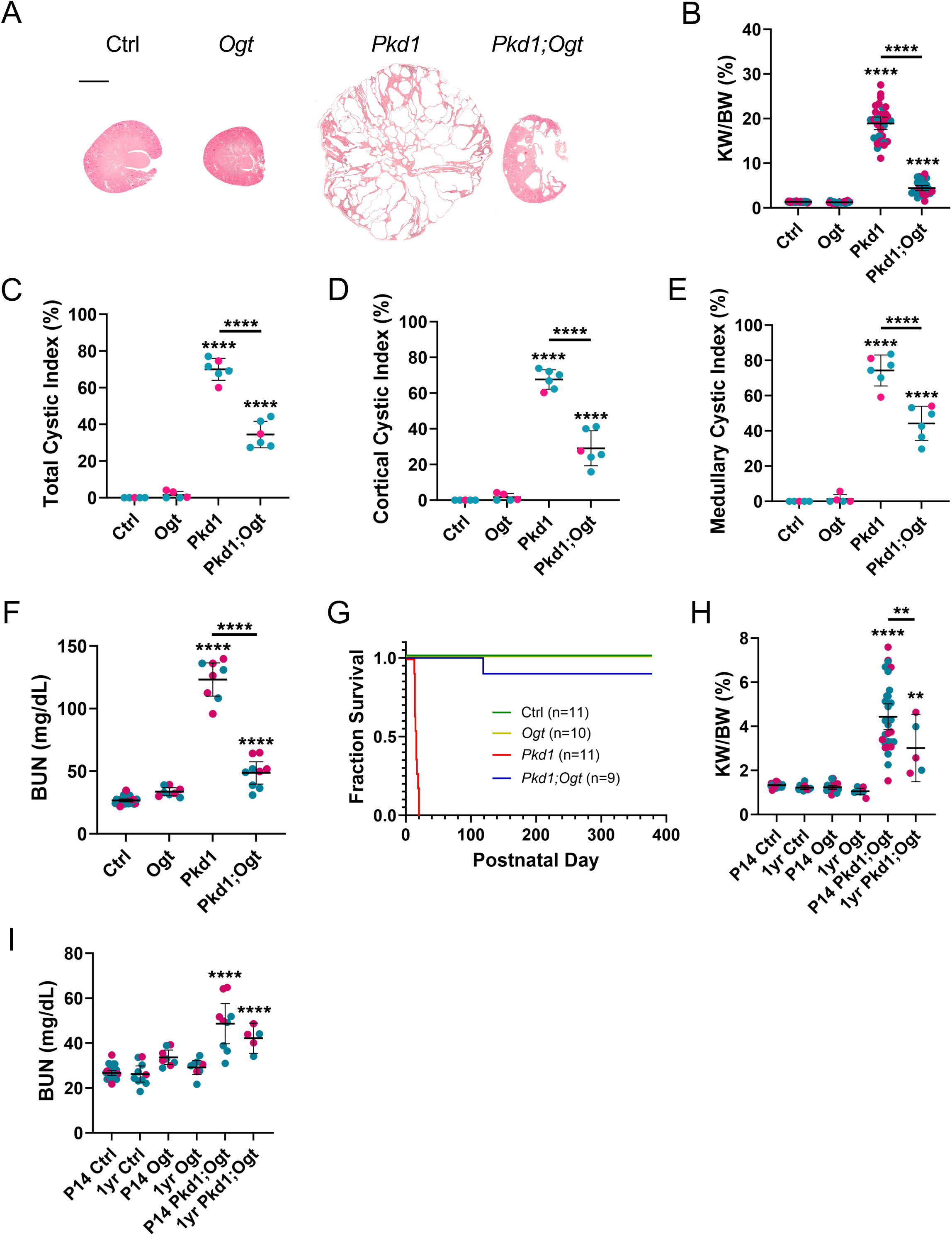
*Ogt* deletion in juvenile *Pkd1* cko kidneys attenuates ADPKD. (A) P14 histology; Scale bar – 1mm (B) kidney weight/body weight (KW/BW) ratios; (C) Percent renal cystogenesis of whole kidney; (D) of cortex; (E) of medulla; (F) blood urea nitrogen (BUN). Blue and pink data points denote male and female mice, respectively. (G) Kaplan Meier Curve. (H) Comparison of KW/BW and (I) BUN levels at P14 and at 1 year of age. *HoxB7-Cre* is expressed in the ureteric bud, which develops into the collecting duct and ureter. At 1 year of age, 3 of the 8 *Pkd1;Ogt* dko mice had hydronephrosis, and thus, were omitted from H and I. Bars represent mean ± SD. Statistical significance was determined by one-way ANOVA followed by Tukey’s test. **p<0.01; ****p<0.0001, compared to control

We examined whether *Ogt* deletion using the HoxB7-Cre recombinase was also beneficial in two additional rapidly progressive ADPKD mouse models. In *Pkd1^RC/fl^* mice, carrying a *Pkd1* floxed allele and the patient mutation, *PKD1* p.R3277C (RC)^29^, *Ogt* deletion reduced KW/BW ratios at P21 from 15.47 ± 3.31 to 6.24 ± 3.91 (mean ± SD; p<0.0001; **Fig S2A**). Additionally, in *Pkd1^RC/dL^* mice, harboring the *RC* allele together with another patient mutation, *PKD1* p.L4132ι1 (ι1L), which deletes a leucine residue in a putative G-protein binding region^30^, *Ogt* deletion reduced KW/BW ratios at P21 from 12.65 ± 3.97 to 8.23 ± 1.15 (p<0.01; **Fig S2B**). The consistent reduction in renal cystogenesis in three rapidly progressive ADPKD mouse models and the extension of mouse survival over 17-fold in *Pkd1;Ogt* dko;*HoxB7-Cre* mice indicates that *Ogt* is a critical gene in ADPKD pathogenesis.

### *Ogt* deletion in a rapidly progressive ADPKD mouse model reduces disease processes

In human and mouse ADPKD kidneys, renal epithelial primary cilia are lengthened, and deletion of certain primary cilia genes in mice attenuates ADPKD severity and reduces primary cilia lengths^28,31^. In *Ogt* cko mice, photoreceptors, which are modified primary cilia, were shortened, and treatment of RPE cells with an irreversible OGT inhibitor also reduced primary cilia lengths^32^. Thus, in addition to regulating metabolism, *Ogt* regulates ciliogenesis. To determine the effect of *Ogt* deletion on kidney cilia lengths, we analyzed cilia of inner cortical collecting duct cells across the different genotypes. We restricted our analyses to the inner cortical collecting duct, since cilia lengths vary along the renal tubule^33^. We stained kidney sections for acetylated α-tubulin, a marker of the ciliary axoneme, and for the *dolichus biflorus agglutinin* (DBA) lectin, which marks the collecting ducts. *Ogt* cko kidneys showed shortened primary cilia. *Pkd1* cko kidneys showed lengthened primary cilia, and *Pkd1;Ogt* dko kidneys showed cilia lengths similar to control (**Figs 2A, 2E**), indicating that deletion of *Ogt* in ADPKD restrains ciliary lengths.

**Figure 2.**
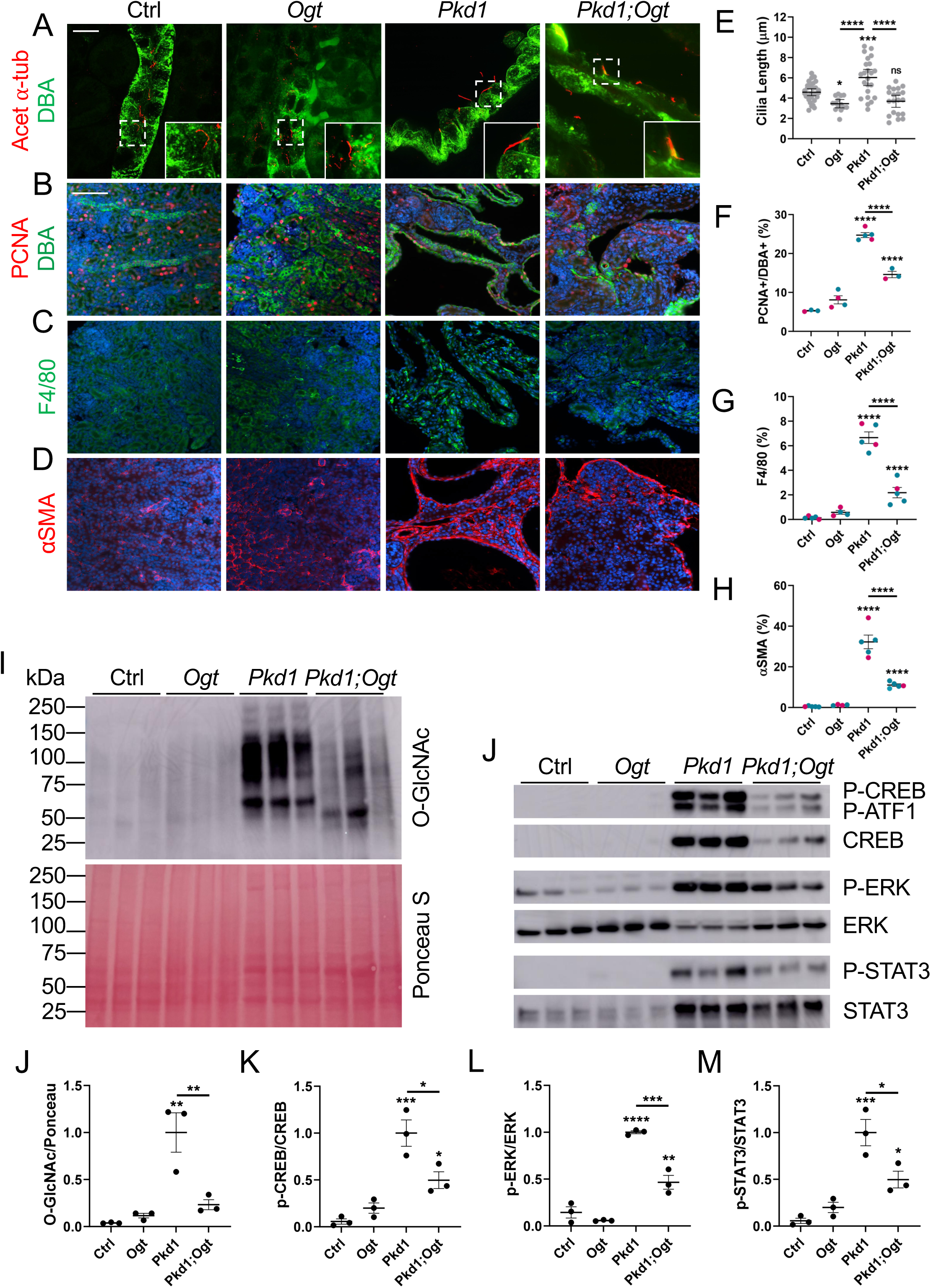
*Ogt* deletion in juvenile *Pkd1* cko kidneys reduces disease hallmarks. Representative images of P14 kidney cortices immunostained for (A) ciliary marker, acetylated α-tubulin (red), with DBA (collecting duct; green); Scale bar - 10μm; (B) cell proliferation marker, PCNA (red) with DBA (green); (C) macrophage marker, F4/80; (D) myofibroblast cell marker, αSMA; with (E, F, G, H) quantification. Scale bar - 50μm. n=5 mice/genotype; 4-5 images were taken of each kidney and quantified. (I) Western blot (WB) analysis on P14 kidney lysates for O-GlcNAc; and for (J) P-STAT3 and STAT3; P-ERK and ERK; and P-CREB and CREB; with (K, L, M, N) quantification; Bars represent mean ± SD. Statistical significance was determined by one-way ANOVA followed by Tukey’s test. *p<0.05; **p<0.01; ***p<0.001; ****p<0.0001, compared to control; ***p<0.001

Increased proliferation of renal tubular epithelia promotes cyst formation, and the progressive growth of kidney cysts causes injury to neighboring parenchyma, resulting in inflammation and fibrosis^34^. We examined the role of *Ogt* deletion on these disease parameters. *Ogt* deletion alone resulted in kidneys that appeared similar to control kidneys, with low proliferation (PCNA+ cells) and few macrophages (F4/80+ cells) and myofibroblasts (α smooth muscle actin+ cells; **Figs 2C-2H, S1G, S1H**). *Pkd1* cko kidneys showed increased proliferation of cyst-lining cells, macrophage infiltration, and increased myofibroblasts. Relative to this, *Pkd1;Ogt* dko kidneys showed reduced proliferation, inflammation and myofibroblasts.

Western blot analyses showed that O-GlcNAc levels were elevated in *Pkd1* cko mouse kidneys and reduced in *Pkd1;Ogt* dko kidneys (**Figs 2I, 2K**). Additionally, activation of CREB, ERK, and STAT3 signaling, which promote renal epithelial fluid secretion, proliferation, and inflammation, were increased in *Pkd1* cko kidneys and reduced in *Pkd1;Ogt* dko kidneys (**Figs 2J-2N**). These data substantiate that deletion of *Ogt* on an ADPKD background attenuates disease severity.

### *Ogt* deletion in a slowly progressive ADPKD mouse model reduces renal cystogenesis and inflammation

We extended our analysis to an adult, slowly progressive ADPKD mouse model. We deleted *Ogt* and *Pkd1* using the *Pax8-rtTA; LC1-Cre* recombinase, which is expressed throughout the renal tubule. Gene deletion was induced from P28-P42, and mice were analyzed at 4 months of age. Deletion of *Ogt* alone largely did not affect kidney morphology, although a few small dilations were occasionally observed (**Figs 3A-3E**). Deletion of *Pkd1* resulted in multiple, large cysts, while deletion of *Ogt* together with *Pkd1* reduced renal cystogenesis and KW/BW ratios. Yet, *Pkd1* cko and *Pkd1;Ogt* dko mice had increased BUN levels (**Fig 3F**). In *Pkd1* cko kidneys, the largest cysts were lined with DBA+ cells, indicating these cysts derived from the collecting duct, while smaller cysts and dilations were lined with Tamms Horsfall Protein positive (THP+) and *Lotus Tetragonolobus* lectin positive (LTL+) cells (**Fig 3G**), marking loop of Henle and proximal tubule origins, respectively.

**Figure 3.**
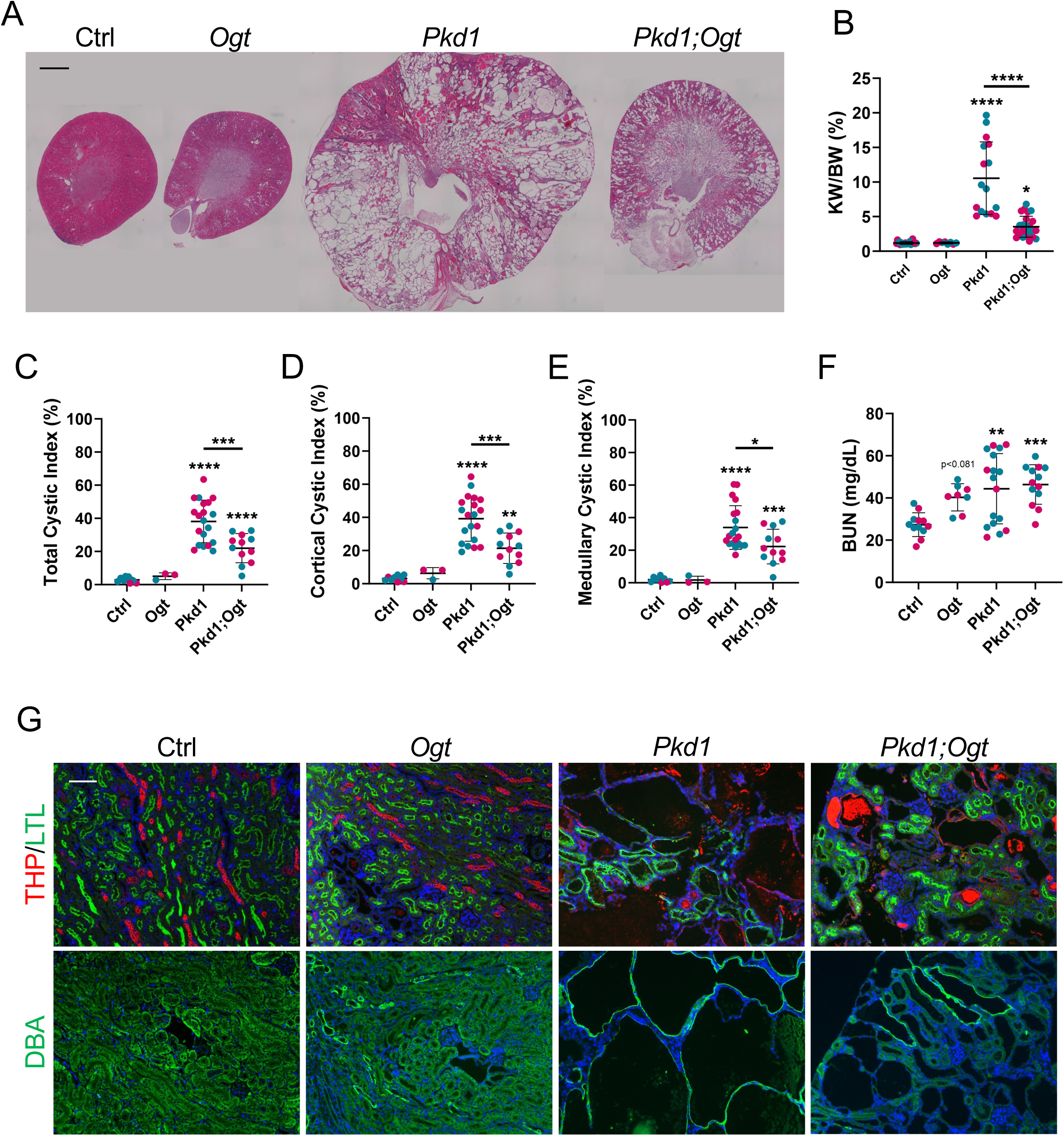
*Ogt* deletion in adult *Pkd1* cko kidneys attenuates ADPKD. (A) Representative images of 4-month-old histology; Scale bar – 1mm (B) KW/BW ratios; (C) Percent renal cystogenesis of whole kidney; (D) cortex; and (E) medulla; (F) BUN. Blue and pink data points denote male and female mice, respectively. Bars represent mean ± SD. Statistical significance was determined by one-way ANOVA followed by Tukey’s test. *p<0.05; **p<0.01; ***p<0.001; ****p<0.0001, compared to control; (G) Staining for proximal tubule (LTL, green) together with loop of Henle (THP, red) and for collecting duct (DBA, green). Scale bar - 100μm

To determine the role of *Ogt* deletion in adult kidney, we analyzed primary cilia of collecting duct cells in the inner cortex across the genotypes. Primary cilia in *Ogt* cko inner cortices were similar in length to those of control, while primary cilia in *Pkd1* cko kidneys were lengthened. In *Pkd1;Ogt* dko kidneys, cilia were longer relative to control, but shortened relative to *Pkd1* cko kidneys. Thus, *Ogt* deletion has capacity to also restrain ciliary lengths in adult ADPKD kidneys (**Figs S3A, S3E**).

To assess the role of *Ogt* deletion on adult ADPKD progression, we examined kidneys for proliferation, inflammation and pro-fibrotic processes. *Ogt* cko kidneys appeared similar to control kidneys, while *Pkd1* cko kidneys showed increased proliferation of cyst-lining cells, macrophage infiltration, and increased myofibroblasts (**Figs S3B-S3H**). Relative to *Pkd1* cko kidneys, *Pkd1;Ogt* dko kidneys showed reduced proliferation, macrophage infiltration and myofibroblasts. O-GlcNAc levels were increased in *Pkd1* cko mouse kidneys and reduced in *Pkd1;Ogt* dko kidneys (**Figs S3I, S3K**). Further, activation of the pro-proliferative and pro-inflammatory ERK and STAT3 signaling pathways was increased in *Pkd1* cko kidney lysates and reduced in *Pkd1;Ogt* dko kidney lysates (**Figs S3J, S3L, S3M**). These data indicate that *Ogt* is a critical molecule also in adult ADPKD pathogenesis.

### OGT inhibition in ADPKD patient-derived primary renal epithelial cells reduces *in vitro* cyst formation

We next examined O-GlcNAcylation in the human disease. Relative to normal human kidney (NHK) sections, O-GlcNAc and OGT, but not OGA, were increased in cyst-lining cells of ADPKD patient kidneys (**Fig 4A**). Furthermore, treatment of ADPKD patient-derived renal epithelial cells with OGT inhibitor, Osmi-1, reduced *in vitro* cyst formation and growth in a dose-dependent manner (**Figs 4B-4D**). Conversely, treatment with OGA inhibitor, Thiamet-G (TMG), increased cyst size. These data indicate that similar O-GlcNAc mechanisms occur in human ADPKD and that downregulation of *Ogt* and O-GlcNAc has therapeutic potential.

**Figure 4.**
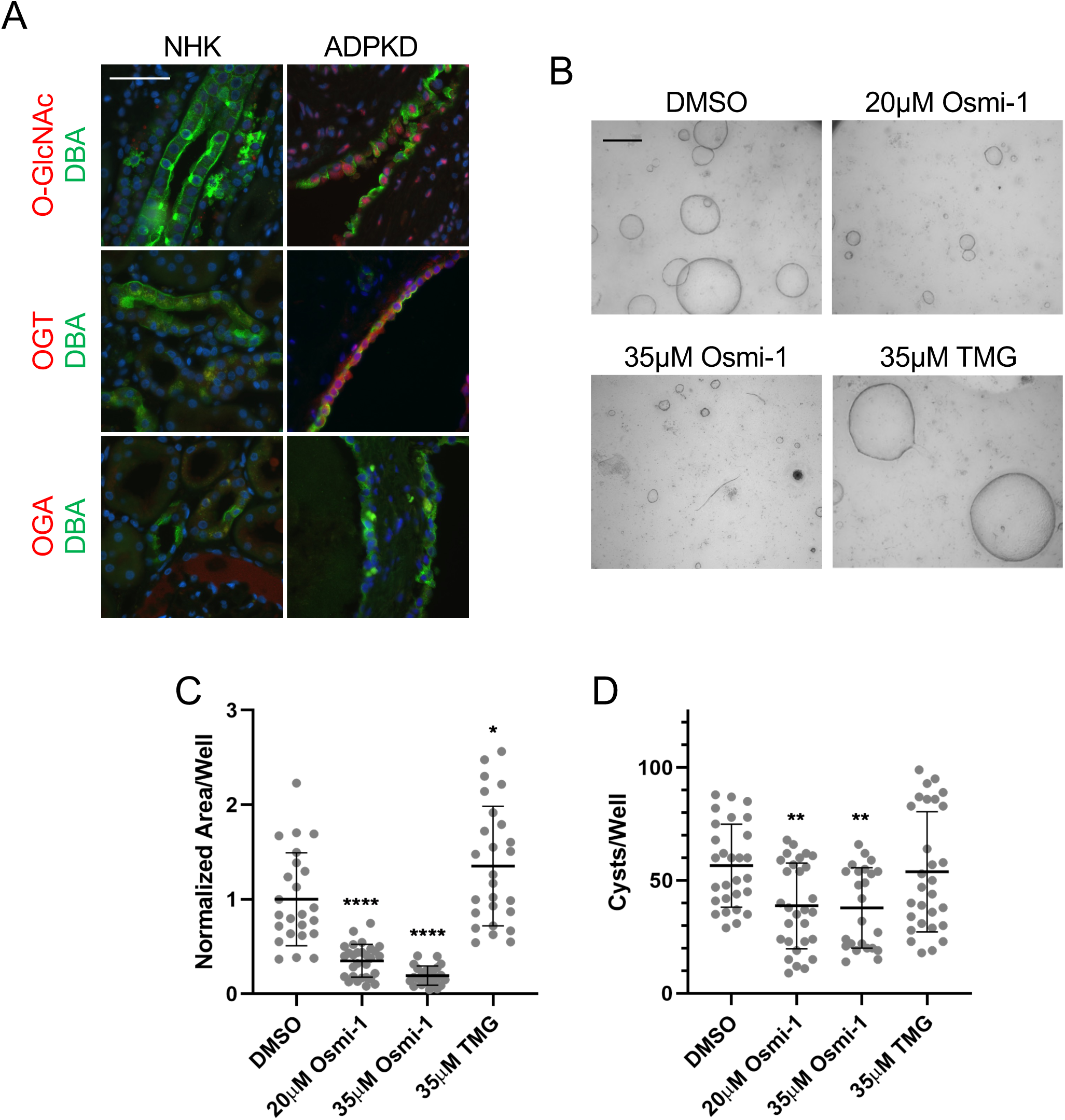
OGT inhibition in human ADPKD cells reduces *in vitro* cyst growth. (A) Representative images of immunostaining for O-GlcNAc (red), OGT (red) and OGA (red) together with collecting duct marker, DBA (green) on NHK (n=5) and ADPKD (n=4) patient kidneys. Samples were derived from male and female individuals, ranging from 44-62 years (Table S3). Scale bar - 50μm (B) Representative micrographs of *in vitro* human ADPKD cysts treated with vehicle (DMSO), Osmi-1 (35μM) or TMG (10μM) in the presence of EGF and FSK to promote cyst formation and growth. Scale bar – 500μm (C) Normalized cyst surface area; (D) cyst number following treatment of ADPKD cells with DMSO, Osmi-1 (20μM, 35μM), or TMG (35μM). Primary renal epithelial cells were derived from male and female ADPKD kidneys (n=5) of individuals ranging from 43-67 years of age (Table S4). Bars represent mean ± SD. Statistical significance was determined using one-way ANOVA followed by Tukey’s test. *p<0.05; **p<0.01; ****p<0.0001

### *Ogt* deletion in juvenile and adult ADPKD mice maintains phosphorylation of AMPK and mitochondrial respiratory chain complex levels

Since OGT is a metabolic sensor^12,13^, we posited that *Ogt* deletion may restrain the metabolic defects that occur in ADPKD. In ADPKD kidneys, activation of the energy sensor, AMPK, is reduced^3,9^. Consistent with this, Western blot analysis showed reduced phosphorylation at Thr183/172 of AMPK in juvenile P14 and adult 4-month-old *Pkd1* cko kidneys (**Figs 5A-5D**). However, relative to *Pkd1* cko kidneys, P-AMPK levels were significantly increased in juvenile and adult *Pkd1,Ogt* dko kidneys. Mitochondrial respiratory chain complex levels are also reduced in ADPKD^35^. In line with this, juvenile and adult *Pkd1* cko kidneys showed diminished levels of all electron transport chain (ETC) complexes. Importantly, in *Pkd1;Ogt* dko kidneys, ETC complex levels were mostly either similar to control levels or significantly increased relative to *Pkd1* cko levels (**Figs 5E-5H; S4**). Thus, deletion of *Ogt* mitigates the reduction in cellular energy sensing and mitochondrial energy production in ADPKD.

**Figure 5.**
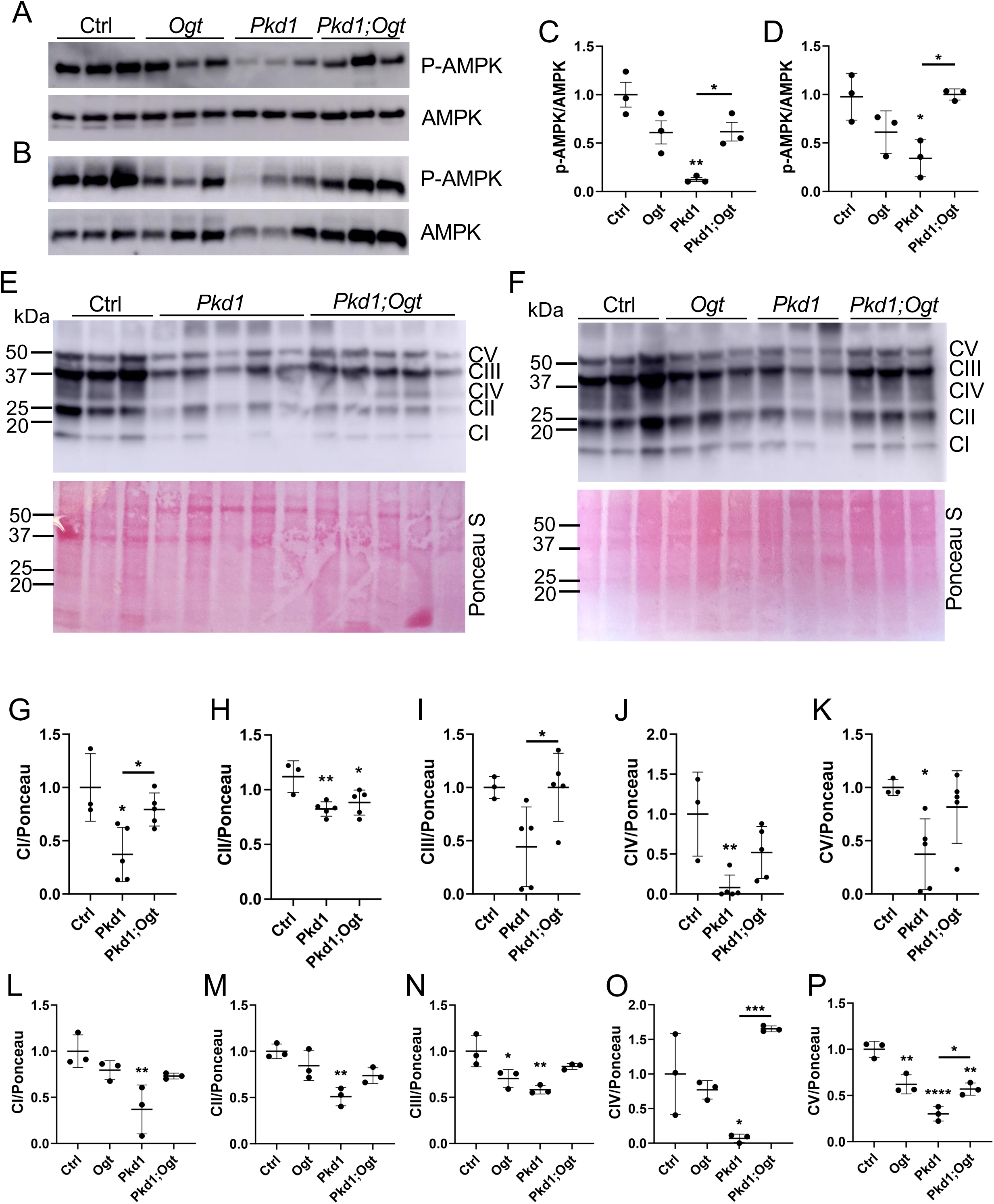
*Ogt* deletion maintains AMPK activation and ETC complex levels (A) WB analysis for P-AMPK and AMPK on P14 *HoxB7-Cre* and (B) 4-month-old *Pax8; LC1-Cre* kidney lysates with (C, D) quantifications, respectively. WB analysis for ETC complexes I-V on (E) P14 *HoxB7-Cre* and (F) 4-month-old *Pax8; LC1-Cre* kidney lysates. (G) Quantification of P14 WB for complex I subunit NDUFB8; (H) complex II subunit SDHB; (I) complex III subunit UQCRCII; (J) complex IV subunit MTCO1; and (K) complex V subunit ATP5A. (L, M, N, O, P) Quantification of 4-month-old WB for complex I-V subunits. Bars represent mean ± SD. Statistical significance was determined by one-way ANOVA followed by Tukey’s test. *p<0.05; **p<0.01; ****p<0.0001, compared to control; ***p<0.001

### *Ogt* deletion in juvenile ADPKD mice restrains metabolic reprogramming

To investigate further the metabolic effects of *Ogt* deletion in *Pkd1* cko kidneys, untargeted metabolomics was performed on P14 control, *Ogt* cko, *Pkd1* cko and *Pkd1;Ogt* dko; *HoxB7-Cre* kidneys. Principle Component Analysis revealed that the 4 genotypes clustered into distinct groups (**Fig 6A**). The control and *Ogt* cko groups clustered more closely together, while the *Pkd1* cko group clustered the furthest away from the control group. The *Pkd1;Ogt* dko group showed slight overlap with the *Pkd1* cko group, but largely remained distinct. In these juvenile mice, sex did not show an effect on clustering.

**Figure 6.**
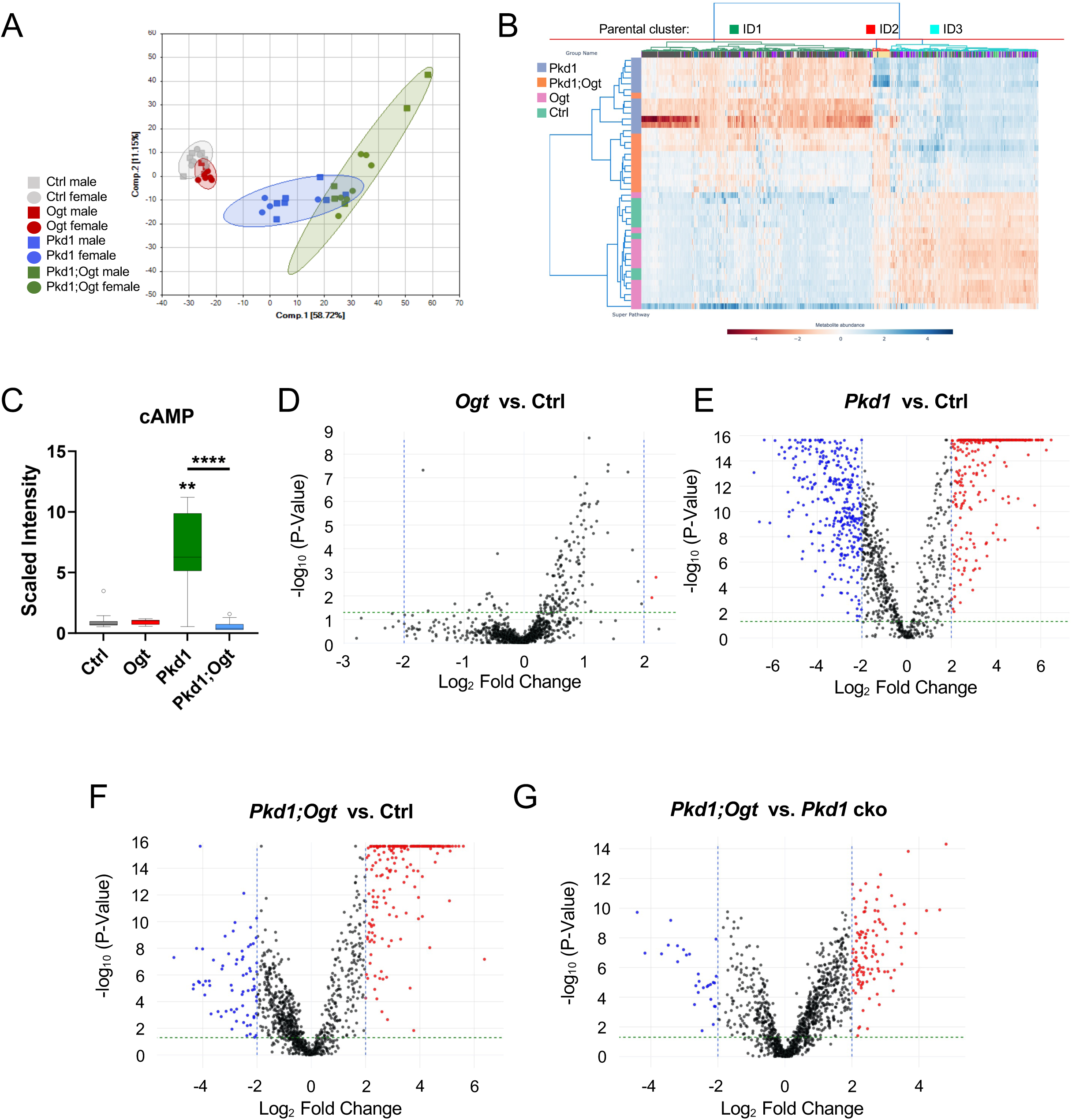
*Ogt* deletion in rapidly progressive *Pkd1* cko kidneys reduces changes in metabolites (A) PCA analysis with ellipses representing 95% confidence intervals; (B) Hierarchical clustering; (C) Box plots of scaled intensity for cAMP. Statistical significance was determined by Kruskal-Wallis test followed by Dunn’s multiple comparisons. **p<0.01, compared to control; ****p<0.0001. (D) Volcano plots of *Ogt* cko vs. control; (E) *Pkd1* cko vs. control; (F) *Pkd1;Ogt* dko vs. control; (G) *Pkd1;Ogt* dko vs. *Pkd1* cko. Data points at -log_10_ p-values >15 represent metabolites that are absent in one group but present in the other group.

Hierarchical clustering also revealed that control and *Ogt* cko groups were most similar, with certain control and *Ogt* cko samples clustering with the *Ogt* cko and control groups, respectively (**Fig 6B**). The *Pkd1* cko group was the most distinct, and the *Pkd1;Ogt* dko group was intermediary between the *Ogt* cko and *Pkd1* cko groups. In the *Pkd1;Ogt* dko group, metabolite levels of parental cluster ID2 were most similar to control and *Ogt* groups, while contrasting with the *Pkd1* cko group. Cluster ID2 was comprised mostly of dipeptides but also included some nucleotide, energy and xenobiotic metabolites. Among these metabolites was cAMP, which promotes cell proliferation and fluid secretion in ADPKD^36^. cAMP was elevated >8-fold in *Pkd1* cko kidneys, but strikingly, was normal in *Pkd1;Ogt* dko kidneys (**Fig 6C**). This suggests a role for *Ogt* in regulating cAMP in ADPKD kidneys.

A total of 1,230 biochemicals – 1,094 known and 136 unknown – were identified. Using a log_2_ fold change of 2 as a cutoff, in *Ogt* cko kidneys relative to control kidneys, only 2 metabolites were identified to be significantly increased, and none were significantly decreased (**Fig 6D**). In contrast, in *Pkd1* cko kidneys, 175 metabolites were increased, and 292 metabolites were decreased (**Fig 6E)**. In *Pkd1;Ogt* dko kidneys, 152 metabolites were increased and 69 metabolites were decreased, relative to control (**Fig 6F**), and 102 metabolites were increased and 28 metabolites were decreased relative to *Pkd1* cko kidneys (**Fig 6G**). Thus, *Ogt* deletion in the ureteric bud and its derivatives has mild effects on kidney metabolism. Moreover, *Ogt* deletion in *Pkd1* cko kidneys halves the number of metabolite changes caused by deletion of *Pkd1*, largely by increasing metabolites that are reduced in *Pkd1* cko kidneys.

In ADPKD, glucose metabolism is dysregulated, resulting in increased glycolysis and increased pentose phosphate pathway (PPP) flux^8^. These pathways are also upregulated in cancer cells, and *Ogt* knockdown suppressed these pathways^37^, suggesting *Ogt* may regulate glucose-dependent pathways. We hypothesized that *Ogt* deletion may similarly rescue the misregulation of glucose-dependent pathways in ADPKD kidneys. In P14 *Pkd1* cko kidneys, glucose and most glycolytic metabolites were reduced, while dihydroxyacetone phosphate (DHAP) was increased (**Figs S5A, S5C-S5G**). In contrast, in *Pkd1; Ogt* dko kidneys, DHAP and most glycolytic metabolite levels were normal or significantly rescued (**Figs S5B, S5C-S5G**). Pyruvate, a glycolytic product, feeds into the TCA cycle. In ADPKD kidneys, intermediates of the first half of the TCA cycle are increased^8^. Consistent with this, in P14 *Pkd1* cko kidneys, the TCA metabolites, citrate, cis-aconitate, isocitrate and α-ketoglutarate, were increased (**Figs S5H, S5I, S6A, S6C, S6D**). However, in *Pkd1;Ogt* dko kidneys, these metabolites were significantly reduced relative to *Pkd1* cko kidneys (**Figs S5H, S5I, S6B-S6D**). In *Pkd1* cko kidneys, fumarate and malate were reduced, but in *Pkd1; Ogt* dko kidneys, these metabolites were significantly increased (**Figs S6E, S6F**). These data suggest that in *Pkd1* cko kidneys, glycolytic metabolites are consumed to enable increased entry of metabolites into the TCA cycle. However, deletion of *Ogt* in *Pkd1* cko kidneys reduced metabolite entry into the TCA cycle, attenuating the dysregulation of glycolysis.

The Hexosamine Biosynthetic Pathway (HBP) is also glucose-dependent^12,13^. In *Pkd1* cko kidneys, most HBP metabolites were reduced (**Figs 7A, 7C-7H**), and O-GlcNAc was elevated (**Figs 2I, 2K**). In contrast, in *Pkd1;Ogt* dko kidneys, most HBP metabolites were normal or significantly rescued (**Figs 7B, 7C-7H**), as were O-GlcNAc levels (**Figs 2I, 2K**). This suggests that in *Pkd1* cko kidneys, HBP metabolites are expended to support increased O-GlcNAcylation, while in *Pkd1;Ogt* dko kidneys, reduced O-GlcNAcylation sustains normal HBP metabolite levels.

**Figure 7.**
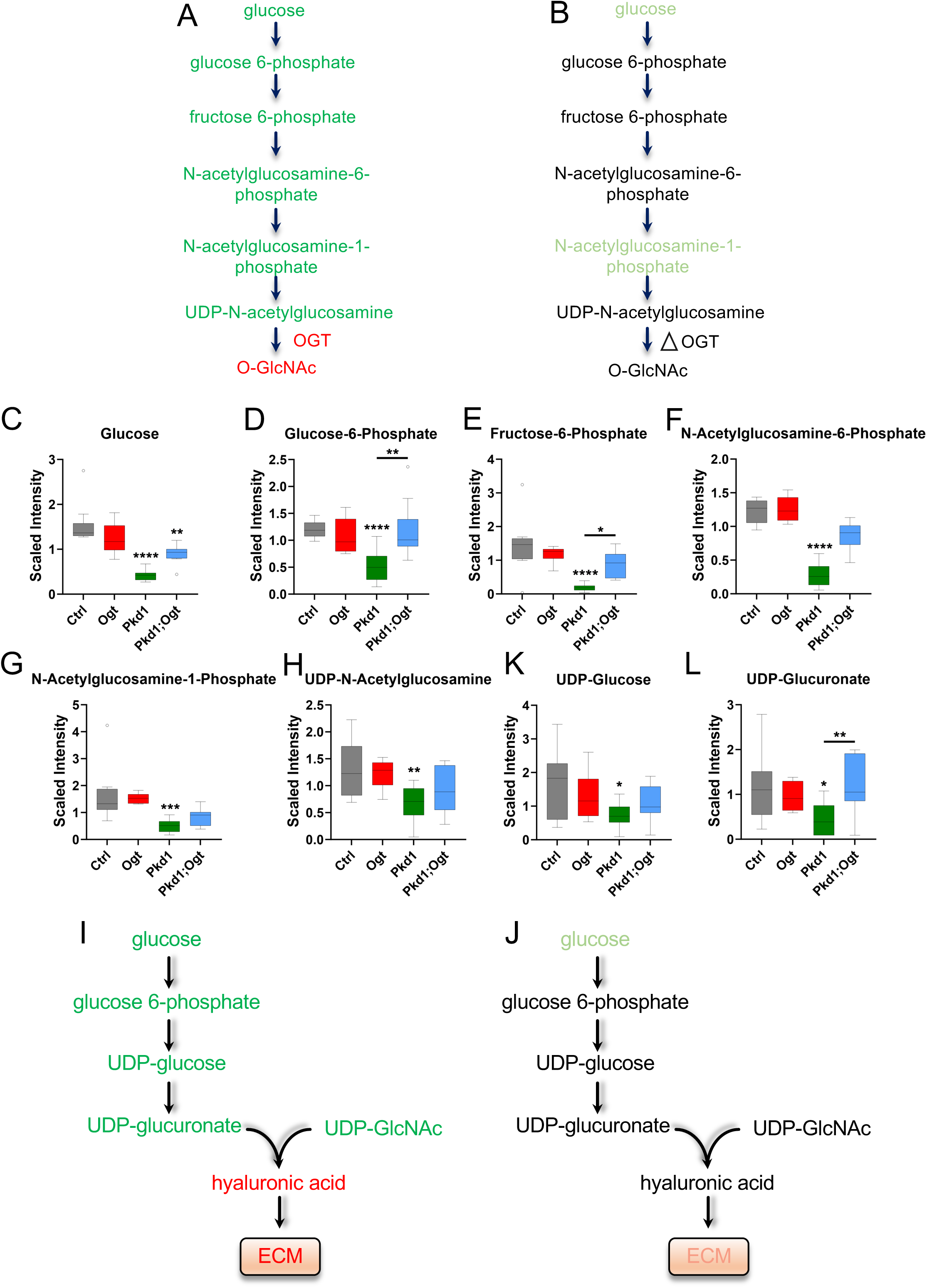
*Ogt* deletion in juvenile *Pkd1* cko kidneys normalizes metabolites of the Hexosamine Biosynthetic Pathway and the Hyaluronic Acid Biosynthesis Pathway. Schematic of HBP metabolite levels for (A) *Pkd1* cko and (B) *Pkd1;Ogt* dko kidneys. Green indicates significantly decreased relative to control. Light green indicates significantly decreased relative to control, but significantly increased relative to *Pkd1* cko. Red indicates significantly increased relative to control. ι1 - deletion (C) Box plots of scaled intensity for glucose; (D) glucose-6-phosphate; (E) fructose-6-phosphate; (F) N-acetylglucosamine-6-phosphate; (G) N-acetylglucosamine-1-phosphate; (H) UDP-N-acetylglucosamine. (I) Schematic of HA biosynthesis pathway metabolite levels for *Pkd1* cko and (J) *Pkd1; Ogt* dko kidneys; (K) Box plots of scaled intensity for UDP-glucose; (L) UDP-glucuronate. Statistical significance was determined using Kruskal-Wallis test followed by Dunn’s multiple comparisons. **p<0.01; ***p<0.001; ****p<0.0001

The hyaluronic acid (HA) biosynthesis pathway is also fueled by glucose. HA is comprised of UDP-GlcNAc and UDP-glucuronate and is a glycosaminoglycan component of the extracellular matrix (ECM)^38^. In normal kidneys, HA is present at low levels and staining in the cortex is not visible (**Fig S7**)^39^. In *Ogt* cko kidney sections, HA biosynthesis pathway metabolites were normal (**Figs 7K, 7L**), and cortical staining of HA was not observed similar to wild-type (**Figs 8A, 8F**). In contrast, in *Pkd1* cko kidneys, metabolites of the HA biosynthesis pathway were decreased (**Figs 7I, 7K, 7L**), and HA was observed around cyst-lining cells and in the interstitial matrix (**Figs 8A, 8F**). In contrast, in *Pkd1;Ogt* dko kidneys, levels of HA biosynthesis metabolites were normal (**Fig 7J, 7K, 7L**), and HA deposition in the ECM was reduced, occurring mostly around blood vessels (**Figs 8A, 8F**).

**Figure 8.**
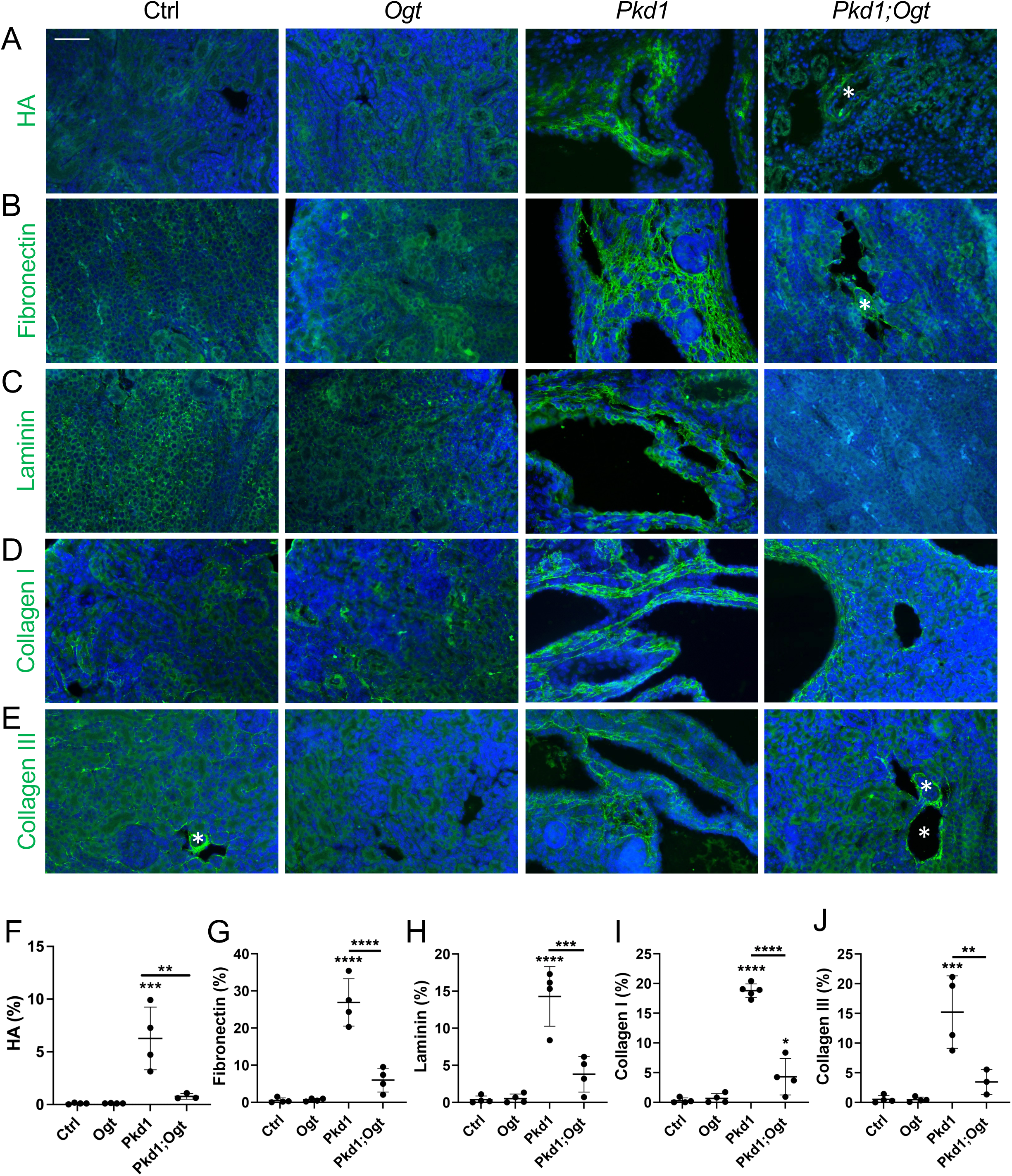
*Ogt* deletion in juvenile *Pkd1* cko kidneys reduces hyaluronic acid and ECM deposition. (A) Cortical HA staining; (B) Immunostaining for cortical fibronectin; (C) laminin; (D) collagen I; (E) collagen III with (F, G, H, I, J) quantification. n=3-5 mice/genotype; 4-5 images were taken of each kidney and quantified. Scale bar - 50μm. White asterisks indicate blood vessels. Bars represent mean ± SD. Statistical significance was determined using one-way ANOVA followed by Tukey’s test. **p<0.01; ***p<0.001; ****p<0.0001

We queried whether alterations of the hyaluronic acid (HA) biosynthesis pathway increasing HA coincided with changes to the ECM. The kidney ECM is comprised of renal epithelial basement membrane and the interstitial matrix^40^. In normal kidneys, the basement membrane is composed of fibronectin, laminin and collagen IV. In wild-type and *Ogt* cko kidney sections, fibronectin and laminin were not easily detectable (**Figs 8B, 8C, 8J, 8H**), reflecting the thinness of the basement membrane. However, in *Pkd1* cko kidneys, fibronectin and laminin were increased around cysts and tubules. Further, collagen I, which is produced by fibroblasts and ADPKD renal tubular epithelial cells^41^, and collagen III, which is produced by fibroblasts, were not visible in wild-type and *Ogt* cko kidneys, but were prominent around cysts and tubules and in the interstitium of *Pkd1* cko kidneys (**Figs 8D, 8E, 8I, 8J**). Notably, in *Pkd1;Ogt* dko kidneys, ECM proteins were markedly reduced. These data suggest that in *Pkd1* cko kidneys, metabolites of the HA biosynthesis pathway were consumed to support increased HA formation, which coincided with increased ECM, while *Ogt* deletion mitigated these alterations. Thus, deletion of *Ogt* in *Pkd1* cko kidneys restrains the dysregulation of glucose-fueled pathways and the intracellular and extracellular pathological processes that ensue.

## Discussion

Our study demonstrates that increased O-GlcNAcylation is pathogenic in ADPKD, presenting O-GlcNAcylation/OGT as a novel metabolic target. Suppressing O-GlcNAc levels in a juvenile mouse model attenuated disease and increased mouse survival from P21 to over a year of age with sustained kidney function. To our knowledge, this is the most effective rescue of a rapidly progressive ADPKD mouse model. Moreover, *Ogt* deletion restrained ADPKD renal cystogenesis in an adult, slowly progressive mouse model, and pharmacological OGT inhibition of patient ADPKD primary renal epithelial cells suppressed *in vitro* cyst growth. This suggests that targeting *Ogt* has therapeutic potential across juvenile and adult ADPKD kidneys and in the human disease.

Metabolomic analysis identified the HBP and the HA biosynthesis pathway as novel pathways in ADPKD, extending the pathogenic effects of altered metabolism to post-translational modification and to the ECM, respectively. HBP flux responds to changes in its nutrient inputs, including glucose, glutamine and acetyl-coA, which are misregulated in ADPKD^8^. Altered O-GlcNAcylation of enzymes can further drive dysregulation of metabolic pathways. In cancer cells, hyper-O-GlcNAcylation of the glycolytic and PPP rate-limiting enzymes, phosphoglycerate kinase 1 (PGK1) and glucose-6-phosphate dehydrogenase (G6PD), increases their enzymatic activities and pathway flux^26,42^. Additionally, O-GlcNAcylation can oppose phosphorylation, altering the cellular post-translational landscape^43^. In cancer cells, hyper-O-GlcNAcylated AMPK reduces phosphorylation and activation of AMPK^44^. Notably, P-AMPK phosphorylates and inhibits the HBP and HA biosynthesis pathway rate-limiting enzymes, glutamine-fructose-6-phosphate amidotransferase (GFAT) and HA synthase 2 (HAS2)^45,46^. Thus, in ADPKD, reduced P-AMPK may decrease GFAT phosphorylation, increasing GFAT activity and culminating in increased UDP-GlcNAc/O-GlcNAcylation. Reduced P-AMPK may also decrease HAS2 phosphorylation, increasing HAS2 activity and HA formation, altering the ECM. Mapping the O-GlcNAcome and phosphoproteome in ADPKD would enhance our understanding of how metabolic dysfunction affects cellular activities and drives disease progression.

The HA biosynthesis pathway represents the first connection between cell metabolism and an extracellular role in ADPKD. While HA is normally low and not visible in healthy kidney cortices^39^, *Pkd1* cko kidneys display increased HA in the ECM. Altered HA metabolism is also evident in human ADPKD, as patients exhibit increased plasma HA^47^. Additionally, TMEM2, a hyaluronidase that breaks down high molecular weight HA into low molecular weight HA, is increased in urinary extracellular vesicles of ADPKD patients^48^. While this degradation is a normal process of HA turnover, chronic sustained breakdown of HA leading to aberrantly elevated levels of low molecular weight HA is pathogenic. In cancer and diabetic nephropathy, increased low molecular weight HA promotes cell proliferation and inflammation^49,50^. These findings suggest HA metabolism warrants further investigation in ADPKD.

Increased hyaluronic acid in the ECM highlights the significant role of the ECM in ADPKD. While cells secrete and continuously remodel the ECM, the ECM in turn provides the metabolic support and physical and mechanical cues that regulate cell behavior and movement^51^. ECM changes occur early in ADPKD kidneys^52^. In early ADPKD stages of the Han:SPRD rat and in cultured human ADPKD cells, ECM changes were associated with increased cell proliferation^41,53,54^. Additionally, type IV collagen mutations identified in patients with Alport Syndrome are now associated with renal cysts, indicating aberrant ECM is causative of renal cysts^55^. ECM changes are also integral to the fibrotic processes that occur in ADPKD^56–60^. By regulating signaling, cell proliferation, tissue morphogenesis, and inflammation, the ECM plays a vital role in early ADPKD renal cystogenesis as well as in fibrosis^57^. Our findings deepen the paradigm that metabolic reprogramming drives ADPKD, exerting not only intracellular but extracellular influences.

In addition to ameliorating these glucose-dependent pathways, *Pkd1;Ogt* dko kidneys maintained normal levels of cAMP, suggesting targeting *Ogt* could be a novel means to restrain cAMP levels. Elucidating this mechanism could lead to an alternative to Tolvaptan, which is the only approved ADPKD therapy. Tolvaptan antagonizes the vasopressin V2 receptor, reducing cAMP levels, but has variable efficacy and causes adverse side effects, such as aquaresis^61^. This underscores the critical need to develop additional therapeutic strategies.

The remediation of multiple metabolic pathways in juvenile *Pkd1;Ogt* dko mice is consistent with the dramatic disease attenuation observed. Despite reduced renal cystogenesis and inflammation in *Pkd1;Ogt* dko; *Pax8;LC1-Cre* mice relative to *Pkd1* cko kidneys, BUN levels were not reduced, unlike in *Pkd1;Ogt* dko; *HoxB7-Cre* mice. *HoxB7* is expressed in the collecting duct, while *Pax8* is expressed throughout the renal tubules, including the proximal tubule. The inability of *Ogt* deletion to reduce BUN levels in *Pkd1* cko; *Pax8;LC1-Cre* mice may indicate that *Ogt* is essential for proximal tubular health. Indeed, in a mouse model of contrast-induced acute kidney injury (AKI) affecting proximal tubules, acute elevation of O-GlcNAcylation prevented AKI^62^. These data indicate that collecting-duct specific targeting of *Ogt* may be required to avoid off-target effects.

A study reported that one-week treatment of juvenile *Pkd1^L^*^3^*^/L^*^3^ mice with OGA inhibitor, TMG, attenuated disease^63^, seemingly contrasting with our genetic findings. However, the relationship between OGT and OGA is complex. While treatment with an OGA inhibitor should theoretically increase O-GlcNAcylation, we have found that one-month TMG treatment of wild-type mice reduced O-GlcNAc levels in the kidneys (data not shown). To understand these results, feedback regulation between OGT and OGA is important to note^64^. Pharmacological OGT inhibition decreases *OGA* expression, and OGA inhibition decreases OGT protein levels. This reflects how critical maintaining O-GlcNAc levels are to preserving cell metabolism and health. Thus, acute versus chronic treatments can yield different results.

Studies in testicular aging, medulloblastoma and infection show that administering OGT inhibitor, Osmi-1, to adult mice effectively reduced O-GlcNAcylation and treated the disease^65–67^, suggesting OGT pharmacological inhibition may be tolerated. However, because acute O-GlcNAcylation is a healthy and necessary process in multiple organs, administering an OGT inhibitor using a cell type-specific drug delivery system, such as a collecting-duct targeting nanoparticle^68^, would be ideal to avoid effects in other cells and organs. Additionally, efforts to develop more specific and potent OGT inhibitors are ongoing^69,70^.

ADPKD patients typically reach renal failure at an average age of 55 years^71^. If a therapy slows ADPKD progression to the extent that patients can reach 90 years of age before developing renal failure, this would constitute a successful intervention. Notably, deletion of *Ogt* in the collecting ducts of juvenile ADPKD mice extended their lifespan of less than 3 weeks to over a year. We posit that collecting duct-specific targeting of O-GlcNAcylation in ADPKD may hold significant therapeutic promise.

## Methods

### Generation of mice

*Pkd1^flox/flox^, Ogt^flox/flox^* and *HoxB7-Cre* mice were obtained from the Jackson Laboratories (Stock numbers 010671, 004860 and 004692, respectively). Mice harboring *Pax8-rtTA* and *LC1-Cre* transgenes, which express the reverse tetracycline-dependent transactivator throughout the renal tubule^72^ and Cre recombinase under the control of an rtTA response element^73^ , respectively, were kindly provided by Dr. Alan Yu, KUMC, with permission from the German Cancer Research Center (DKFZ). *Pkd1^RC/RC^* mice were a generous gift from Dr. Peter Harris, Mayo Clinic^29^. *Pkd1^τιL/+^* mice were generated as described^30^. *Pkd1^flox/flox^;Ogt^flox/+^*or *Pkd1^flox/+^;Ogt^flox/+^* females and *Pkd1^flox/flox^;Ogt^flox/y^*, *HoxB7-Cre* males were generated and mated to produce *Pkd1* cko, *Ogt* cko and *Pkd1;Ogt* dko; *Hoxb7-Cre* mice. *Pkd1^flox/+^;Ogt^flox/+^* females and *Pkd1^RC/+^;Ogt^flox/y^*; *HoxB7-Cre* males were generated and mated to produce *Pkd1^RC/flox^*; *HoxB7-Cre* mice, with or without *Ogt* deletion. Similarly, *Pkd1^τιL/+^;Ogt^flox/+^*females and *Pkd1^RC/+^;Ogt^flox/y^*; *HoxB7-Cre* males were generated and mated to produce *Pkd1^RC/τιL^* mice with or without *Ogt* deletion. Finally, *Pkd1^flox/flox^;Ogt^flox/+^; Pax8-rtTA+; LC1-Cre+* or *Pkd1^flox/+^;Ogt^flox/+^;Pax8-rtTA+; LC1-Cre+* females and *Pkd1^flox/flox^;Ogt^f/y^*,*Pax8-rtTA+;LC1-Cre+* males were generated. These parental lines were mated to produce single and double knock-out (dko) mice. To induce *Pkd1* and/or *Ogt* deletion in adult-onset models, doxycycline (2μg/g; ThermoScientific, J60579.22) and 3% glucose was added to the drinking water from P28-P42. All mouse lines were maintained on a pure C57BL6/J background (backcrossed ≥10 generations). Animal procedures were conducted in accordance with KUMC-IACUC and AAALAC rules and regulations.

### Histology, immunofluorescence and hyaluronic acid binding protein assay

Paraffin-embedded kidney tissue sections (7 μm) were deparaffinized, rehydrated through an ethanol series, then stained with hematoxylin and eosin. Alternatively, once deparaffinized and rehydrated, tissue sections were subjected to antigen retrieval, by steaming for 25 minutes in Sodium Citrate Buffer (10 mM Sodium Citrate, 0.05% Tween 20, pH 6.0), returned to room temperature, then rinsed 10 times in distilled water. As an additional step to the antigen retrieval, sections were incubated with 1% SDS in PBS for 5 minutes, then washed in PBS. Immunofluorescence was performed as described^74^, using primary antibodies alone or together with LTL or DBA lectins (Supplementary Table S1) followed by secondary antibodies (Supplementary Table S2). Sections were mounted with DAPI Fluoromount-G (Electron Microscopy Sciences, 17984-24), ProLong Gold (Invitrogen, P10144) or ProLong Gold with DAPI (Invitrogen, P36935).

To detect HA, HA binding protein assays were carried out as described^75^. Tissue sections were deparaffinized and rehydrated in PBS, then incubated sequentially in avidin block, biotin block (Vector Labs, SP-2001) and goat serum (Fisher Scientific, 16-210-064). Sections were treated with saline or hyaluronidase (Sigma-Aldrich, H3884) at 37^0^C for 1 hour, which served as a negative control. Sections were washed in PBS and incubated with biotinylated hyaluronic acid binding protein (EMD-Millipore-Calbiochem, 38599) overnight at 4^0^C. Sections were washed in PBS, incubated with ABC reagent (Vector Labs, PK-6101), washed in PBS, then incubated with TSA reagent plus fluorescein (Perkin Elmer, NEL741001KT). Sections were washed in PBS, then mounted in DAPI Fluoromount-G (Electron Microscopy Sciences, 17984-24). Staining was imaged using a Nikon 80i microscope with a photometrics camera or a Leica DMLB microscope with an Infinity5 Lumenera camera. Immunostaining for cilia was imaged using a TI2-E inverted microscope attached to CSU-W1 Spinning-disk confocal with SoRa super-resolution.

IF images were quantified using ImageJ. To measure cilia length, the line tool was used to trace the scale bar of an image and determine how many pixels equals a micron. Subsequently, cilia lengths were measured using the line tool and length measurements converted from pixels into microns. To quantify area of other fluorescent stains, single channel fluorescence images were converted to a 16-bit image (Image ®Type ® 16-bit). The threshold tool was used to measure the area of a specific stain (Image ® Adjust ® Threshold). The threshold was then adjusted to measure the whole kidney area. The area of the specific stain was divided by the area of the kidney tissue section.

### Western blot

Western blots were performed as described^76^. Briefly, kidney pieces were homogenized in RIPA Buffer, pH 7.6 containing GlcNAc (1% NP-40, 10mM Tris, 1mM DTT, 2mM EDTA, 150mM NaCl, 0.5% deoxycholic acid, sodium salt, 40mM GlcNAc, 0.1% SDS) and protease inhibitors (Thermofisher, A32965) using a BulletBlender Storm 24 (NextAdvance). To determine protein concentrations, BCA assays (ThermoFisher, 23227) were performed according to manufacturer’s instructions. Protein lysates (50μg) were boiled for 5 minutes for most primary antibodies or heated at 37^0^C for 5 minutes for the mouse OxPhos antibody cocktail. Lysates were run on a gel, then transferred onto a Polyvinylidene Fluoride (PVDF) membrane (Sigma, P2938). Membranes were stained with Ponceau S (Sigma, P3504), then incubated with primary and secondary antibodies (Supplementary Tables S1 and S2).

Western blots were quantified using ImageJ software. Western blot and Ponceau images were converted to 16-bit (Image ®Type ® 16-bit). Band(s) within the lane were selected using the rectangle tool (Analyze ® Gels ® Select First Lane). The rectangle was dragged to select the band(s) in the next lane (Analyze ® Gels ® Select Next Lane), and subsequently, all other lanes. Selections were plotted (Analyze ® Gels ® Plot Lanes). On the plots, lines were drawn to demarcate the base of a peak, and the wand tool was used to quantify area under the peak. Three to five bands on a Ponceau image were quantified and summed to provide the relative value of total protein within a lane. Value of the band of interest on a Western blot was divided by the value of the Ponceau bands.

### Blood Urea Nitrogen Measurements

Blood was retrieved either through cardiac puncture of anesthetized mice or via draining of trunk blood of euthanized mice. Blood was collected in Microvette CB 300 Blood Collection System tubes (Kent Scientific, KMIC-SER), then centrifuged at 2000g at room temperature for 10 minutes to collect serum. BUN levels were measured using the QuantiChrom Urea Assay Kit (BioAssay Systems, DIUR-100) according to manufacturer’s protocol.

### ADPKD and normal human kidney (NHK) sections

For immunofluorescence, ADPKD (K472, K485, K488, K493) and NHK (K465, K482, K484, K491, K494) tissue sections (Supplementary Table S3) were obtained from the PKD Biomarkers, Biomaterials, and Cellular Models Core of the Kansas PKD Center. The protocol for the use of discarded human tissues complied with federal regulations and was approved by the Institutional Review Board at KUMC^76^.

### In vitro cyst formation

ADPKD cells (K387, K419, K441, K443, K488; Supplementary Table S4) were subjected to an *in vitro* cyst assay as described^76^. Briefly, cells (4,000 cells/well) were dispersed in cold Type I collagen (Advanced Biomatrix; San Diego, CA) in wells of a 96-well plate, warmed to 37 °C to allow for the collagen gel to polymerize. Defined media (DMEM/F12, Pen/Strep, ITS Culture Supplement (Fisher), 5 × 10^−8^ M Hydrocortisone and 5 × 10^−12^ M Triiodothyronine) supplemented with forskolin (FSK, 5 μM) and EGF (5 ng/ml), was placed onto collagen-suspended cells, and refreshed every 2-3 days over a 7-day period to stimulate *in vitro* cyst formation. Once microcysts were observed, cultures were fed with media containing either DMSO (Sigma), 20 μM Osmi-1 (MedChemExpress, 119738), or 35μM TMG (Selleckchem, S7213). Following the microcyst assay, culture gels were fixed in 0.5% paraformaldehyde, and microcysts were photographed with a digital camera attached to an inverted microscope Nikon Eclipse TE2000-U, objective (2X). Diameters of spherical cysts with distinct lumens were measured using Image-Pro Premier 9.2. Diameters of treatment groups were normalized to the mean diameter of the DMSO control group. Assays were performed in six replicate wells per biological replicate.

### Statistics

GraphPad Prism 10 software was used to perform statistical analyses. An unpaired t-test was used to determine statistical significance (p<0.05) between two groups. ANOVA or Kruskal-Wallis tests were used for more than two groups, with or without normal distributions, respectively.

### Metabolomics

Untargeted metabolomic analysis was performed on control (n=12), *Ogt* cko (n=8), *Pkd1* cko (n=12), *Pkd1; Ogt* dko; *HoxB7-Cre* (n=11) kidneys of P14 male and female mice. Mice were injected i.p. with 60mg/kg pentobarbital. Once mice were unresponsive to a toe pinch about 5-7 minutes following the injection, the peritoneal cavity was opened, the renal artery was clamped, and the left kidney was extracted, weighed and snap-frozen in liquid nitrogen between two pre-frozen metal plates. Kidneys were extracted, weighed and snap-frozen in less than 30 seconds. Renal tissue (≥50mg) was sent to Metabolon, Inc.

At Metabolon, Inc., metabolites of frozen kidney tissues were extracted in cold organic solvent and analyzed using liquid chromatography tandem mass spectrometry (LC/MS-MS) as described^77^. An automated liquid handling robot (Hamilton LabStar, Hamilton Robotics) and GenoGrinder (Glen Mills, 2000) were used to extract and mix the samples, from which 4 aliquots were made. Samples were run on 4 metabolomic platforms, representing 4 separate LC–MS/MS arms: 1) positive ionization chromatographically optimized for hydrophilic compounds (LC–MS/MS Pos Polar); 2) positive ionization chromatographically optimized for hydrophobic compounds (LC–MS/MS Pos Lipid); 3) negative ionization optimized conditions (LC–MS/MS Neg); and 4) negative ionization with Hydrophilic Interaction Liquid Chromatography (HILIC; LC–MS/MS Polar). An Acquity Ultra Performance Liquid Chromatography (UPLC; Waters) system at 40–50^0^C was used to separate metabolites in samples. Q Exactive high resolution/accurate mass spectrometers with heated electrospray ionization (HESI-II) sources and Orbitrap mass analyzers (ThermoFisher Scientific) operated at 35,000 mass resolution were used. Data acquisition alternated between full scan MS and data-dependent MS^n^ scans. The scan range generally covered 70–1000 m/z.

Biochemicals were normalized to the mass of the sample extracted, then log transformed. Two-way ANOVA was performed to identify biochemicals exhibiting significant interaction and main effects for genotype and sex.

## Acknowledgements

We thank members of the Jared Grantham Kidney Institute for helpful discussions. We also thank Jing Huang of the KUMC Histology Core, which is supported by NIH U54HD090216. This work was further supported by an ASN KidneyCure Fellowship to MAK; K-INBRE summer scholar awards to NM and JJ (P20 GM103418); PKD summer studentships to RA, CB and CR (U54 DK126126); Jared Grantham Pilot and Feasibility grants to CS and PVT; a DoD Discovery Award (PR211628), a PKD Foundation research grant and a KUMC Lied Preclinical Grant to PVT. ADPKD tissues and cells were provided by the Kansas Polycystic Kidney Disease Research and Translation Core Center of the PKD-RRC Consortium (U54 DK126126).

---

**Supplementary Figure S1.** *Ogt* deletion in rapidly progressive *Pkd1* cko; *HoxB7-Cre* mice reduces renal cystogenesis at P21. (A) Histology; (B) KW/BW; (C) Renal cystogenesis of whole kidney (D) cortex (E) medulla; (F) Quantification of immunostaining for F4/80 and (G) αSMA. While most *Pkd1* cko mice survived until P14, very few survived until P21, accounting for the low n of P21 *Pkd1* cko mice. Bars represent mean ± SD. Statistical significance was determined by one-way ANOVA followed by Tukey’s test. ****p<0.0001, compared to control

**Supplementary Figure S2.** *Ogt* deletion in rapidly progressive *Pkd1^RC/fl^*and *Pkd1^RC//τιL^* mouse models reduces kidney weight. (A) %KW/BW at P21 of *Pkd1^RC/fl^* and (B) *Pkd1^RC//τιL^* mice with and without *Ogt* deletion using the HoxB7-Cre recombinase. Bars represent mean ± SD. Statistical significance was determined using an unpaired t-test. **p<0.01; ****p<0.0001

**Supplementary Figure S3.** *Ogt* deletion in adult *Pkd1* cko kidneys reduces disease hallmarks. Representative images of 4-month-old kidney cortices immunostained for (A) ciliary marker, acetylated α-tubulin (red), with DBA (collecting duct; green); Scale bar - 10μm (B) PCNA (red) with DBA (green); (C) F4/80 (D) αSMA; Scale bar - 50μm and (E, F, G, H) quantification; (I) Western blot analysis on 4-month-old kidney lysates for O-GlcNAc; (J) P-ERK and ERK; P-STAT3 and STAT3; and (L, M, N) Quantification; Bars represent mean ± SD. Statistical significance was determined by one-way ANOVA followed by Tukey’s test. *p<0.05; ****p<0.0001, compared to control

**Supplementary Figure S4.** ETC complex levels in juvenile *Ogt* cko kidneys (A) WB analysis for ETC complexes on P14 control and *Ogt* cko;*HoxB7-Cre* kidney lysates with (B, C, D, E, F) complex I-V quantification. Bars represent mean ± SD. Statistical significance was determined by unpaired t-test. *p<0.05

**Supplementary Figure S5.** *Ogt* deletion in rapidly progressive *Pkd1* cko kidneys reduces changes in glycolytic metabolites. Schematic of glycolytic metabolite levels for (A) *Pkd1* cko and (B) *Pkd1; Ogt* dko kidneys; Green indicates significantly decreased relative to control. Light green indicates significantly decreased relative to control, but significantly increased relative to *Pkd1* cko. Red indicates significantly increased relative to control. (C) Box plots of scaled intensity for DHAP; (D) 3-phosphoglycerate; (E) 2-phosphoglycerate; (F) phosphoenolpyruvate; (G) pyruvate; (H) citrate; (I) aconitate. Statistical significance was determined by Kruskal-Wallis test followed by Dunn’s multiple comparisons.. *p<0.05; **p<0.01; ***p<0.001; ****p<0.0001, compared to control

**Supplementary Figure S6.** *Ogt* deletion in rapidly progressive *Pkd1* cko kidneys reduces changes in metabolites of the TCA cycle. Schematic of TCA metabolite levels for (A) *Pkd1* cko and (B) *Pkd1; Ogt* dko kidneys; Green indicates significantly decreased relative to control. Light green indicates significantly decreased relative to control, but significantly increased relative to *Pkd1* cko. Red indicates significantly increased relative to control. (C) Box plots of scaled intensity for isocitrate; (D) α-ketoglutarate; (E) fumarate; (F) malate. Statistical significance was determined by Kruskal-Wallis test followed by Dunn’s multiple comparisons. *p<0.05; **p<0.01; ****p<0.0001, compared to control

**Supplementary Figure S7.** HA in control kidney. (A) HA in P14 control kidney showing cortex and medulla on right side of image. Bright green spots in interstitium of medulla indicate HA. Proximal tubules on left side of image show increased green intensity due to nonspecific uptake of fluorescent stain. Scale bar - 100μm. (B) HA in P14 control medulla.

**Supplementary Table S1.** Primary antibodies used in immunofluorescence (IF) and Western blot (WB)

**Supplementary Table S2.** Secondary antibodies used in immunofluorescence (IF) and Western blot (WB)

**Supplementary Table S3.** NHK and ADKPD samples used in IF

**Supplementary Table S4.** ADPKD samples used in *in vitro* cyst assays

## Notes

### Competing Interest Statement

The authors have declared no competing interest.

