## Supplemental tables and figures for "Targeting *Ogt* in ADPKD mitigates metabolic reprogramming and renal cystogenesis, extending survival"

M. Kavanaugh *et al.*

**Supplementary Tables S1-S4**

**Table S1.** Primary antibodies used in immunofluorescence (IF) and Western blot (WB)

| Primary antibody or lectin | Catalog no. | Dilution | Supplier | Experiment |
|---|---|---|---|---|
| **acetylated α-tubulin** | T7451 | 1:4000 | Sigma-Aldrich Corp. | IF |
| **αSMA** | ab5694 | 1:500 | Abcam | IF |
| **F4/80** | 30325S | 1:400 | Cell Signaling Technology | IF |
| **PCNA** | 13110S | 1:300 | Cell Signaling Technology | IF |
| **THP** | sc-271022 | 1:100 | Santa Cruz Biotechnology | IF |
| **DBA** | FL-1031 | 1:100 | Vector Laboratories | IF |
| **LTL** | FL-1321 | 1:300 | Vector Laboratories | IF |
| **O-GlcNAc** | MA1-072 | 1:100 | ThermoFisher | IF and WB |
| **OGT** | | 1:200 | Kind gift from Dr. G. Hart | IF |
| **OGA** | | 1:2000 | Kind gift from Dr. G. Hart | IF |
| **fibronectin** | F3648 | 1:400 | Sigma-Aldrich Corp. | IF |
| **laminin** | L9393 | 1:100 | Sigma-Aldrich Corp. | IF |
| **collagen I** | 1310-01 | 1:50 | Southern Biotech | IF |
| **collagen III** | 1330-01 | 1:50 | Southern Biotech | IF |
| **P-ERK** | 4370S | 1:1000 | Cell Signaling Technology | WB |
| **ERK** | 4696S | 1:1000 | Cell Signaling Technology | WB |
| **P-STAT3** | 9145S | 1:1000 | Cell Signaling Technology | WB |
| **STAT3** | 9139S | 1:1000 | Cell Signaling Technology | WB |
| **P-CREB** | 9198 | 1:1000 | Cell Signaling Technology | WB |

| | | | | |
|---|---|---|---|---|
| **CREB** | 9197 | 1:1000 | Cell Signaling Technology | WB |
| **Total OxPhos Rodent WB Antibody Cocktail** | AB110413 | 1:4000 | Abcam | WB |

**Table S2.** Secondary antibodies used in immunofluorescence (IF) and Western blot (WB).

| Secondary antibody | Catalog no. | Dilution | Supplier | Experiment |
|---|---|---|---|---|
| **rabbit, Alexa Fluor 488** | A11034 | 1:500 | ThermoFisher Scientific | IF |
| **mouse, Alexa Fluor 488** | A11001 | 1:500 | ThermoFisher Scientific | IF |
| **goat, Alexa Fluor 488** | A21467 | 1:500 | ThermoFisher Scientific | IF |
| **rabbit, Alexa Fluor 594** | A11012 | 1:500 | ThermoFisher Scientific | IF |
| **mouse, Alexa Fluor 594** | A11005 | 1:500 | ThermoFisher Scientific | IF |
| **rabbit, HRP-conjugated** | 7074S | 1:5000 | Cell Signaling Technology | WB |
| **mouse, HRP-conjugated** | 7076S | 1:5000<br>1:10,000 for O-GlcNAc<br>1:15,000 for OxPhos | Cell Signaling Technology | WB |

**Table S3.** NHK and ADPKD samples used for IF

| Sample ID | Sex | Age (years) |
|---|---|---|
| **NHK K465** | female | 55 |
| **NHK K482** | female | 48 |
| **NHK K484** | male | 44 |
| **NHK K491** | female | 62 |
| **NHK K494** | female | 46 |
| **ADPKD K472** | female | 52 |
| **ADPKD K485** | male | 54 |
| **ADPKD K488** | female | 53 |
| **ADPKD K493** | female | 49 |

**Table S4.** ADPKD samples used for *in vitro* cyst assays

| Sample ID | Sex | Age (years) |
|---|---|---|
| **ADPKD K319** | male | 48 |
| **ADPKD K419** | female | 43 |
| **ADPKD K441** | male | 60 |
| **ADPKD K443** | female | 67 |
| **ADPKD K488** | female | 53 |

Figure S1

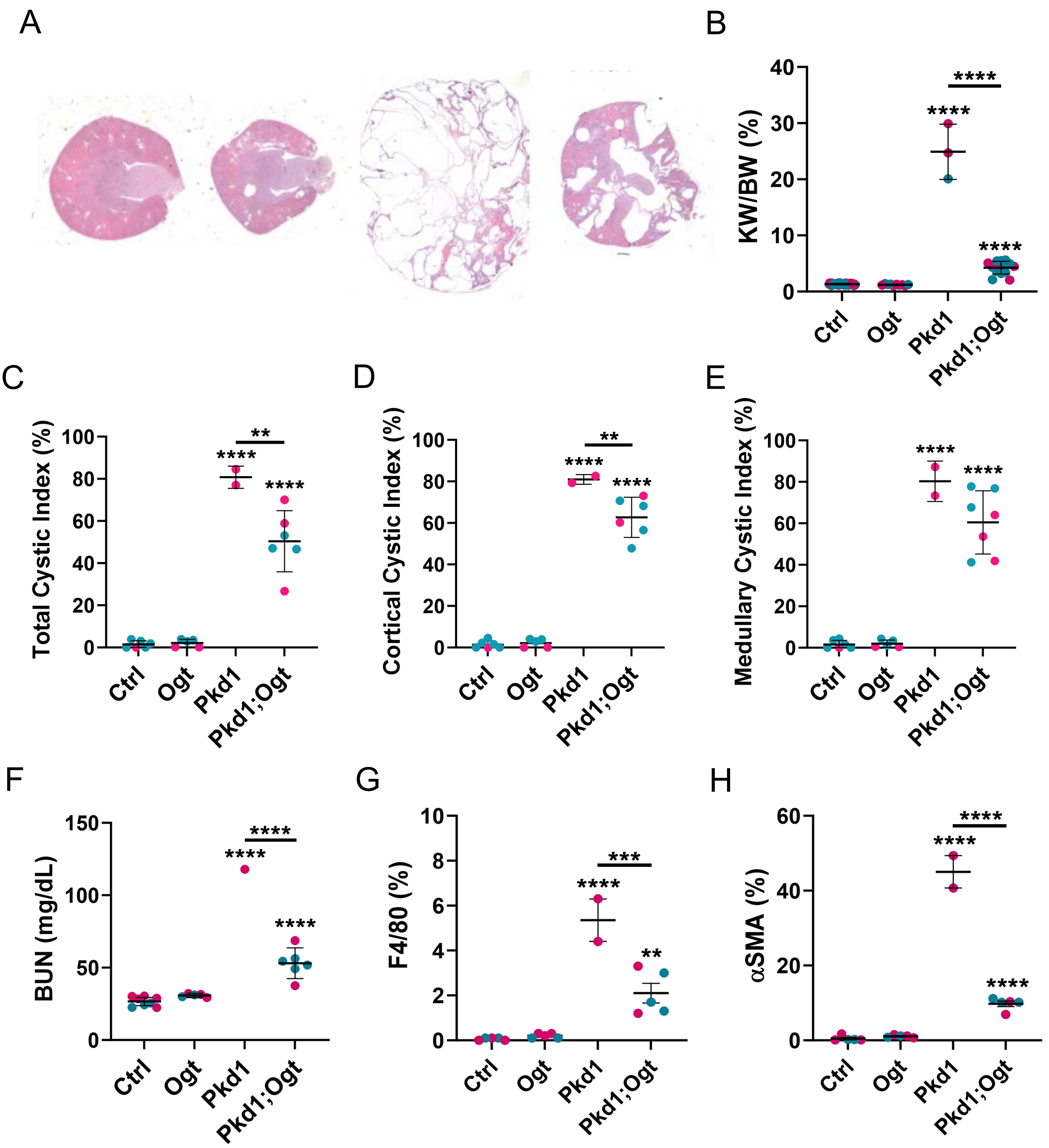

A
B
KW/BW (%)
40
30
20
10
0
****
****
****
Ctrl
Ogt
Pkd1
Pkd1;Ogt
C
Total Cystic Index (%)
100
80
60
40
20
0
**
****
****
D
Cortical Cystic Index (%)
E
Medullary Cystic Index (%)
F
BUN (mg/dL)
150
100
50
0
G
F4/80 (%)
10
8
6
4
2
0
***
**
H
αSMA (%)
60
40
20
0

Figure S2

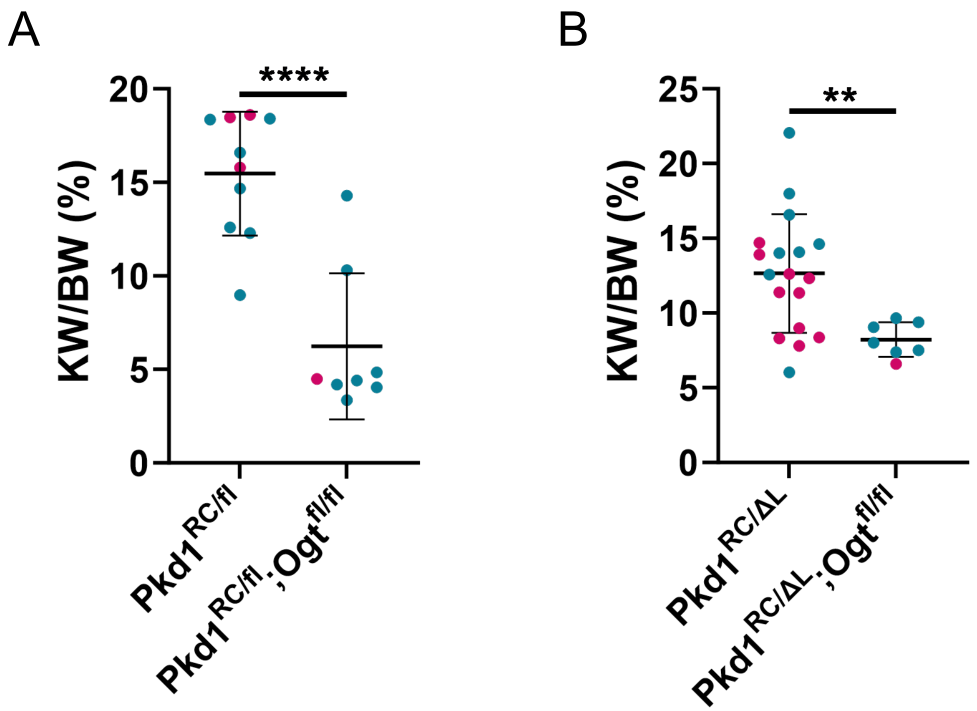
A
B
KW/BW (%)
****
**
0
5
10
15
20
25
Pkd1RC/fl
Pkd1RC/fl;Ogtfl/fl
Pkd1RC/ΔL
Pkd1RC/ΔL;Ogtfl/fl

Figure S3

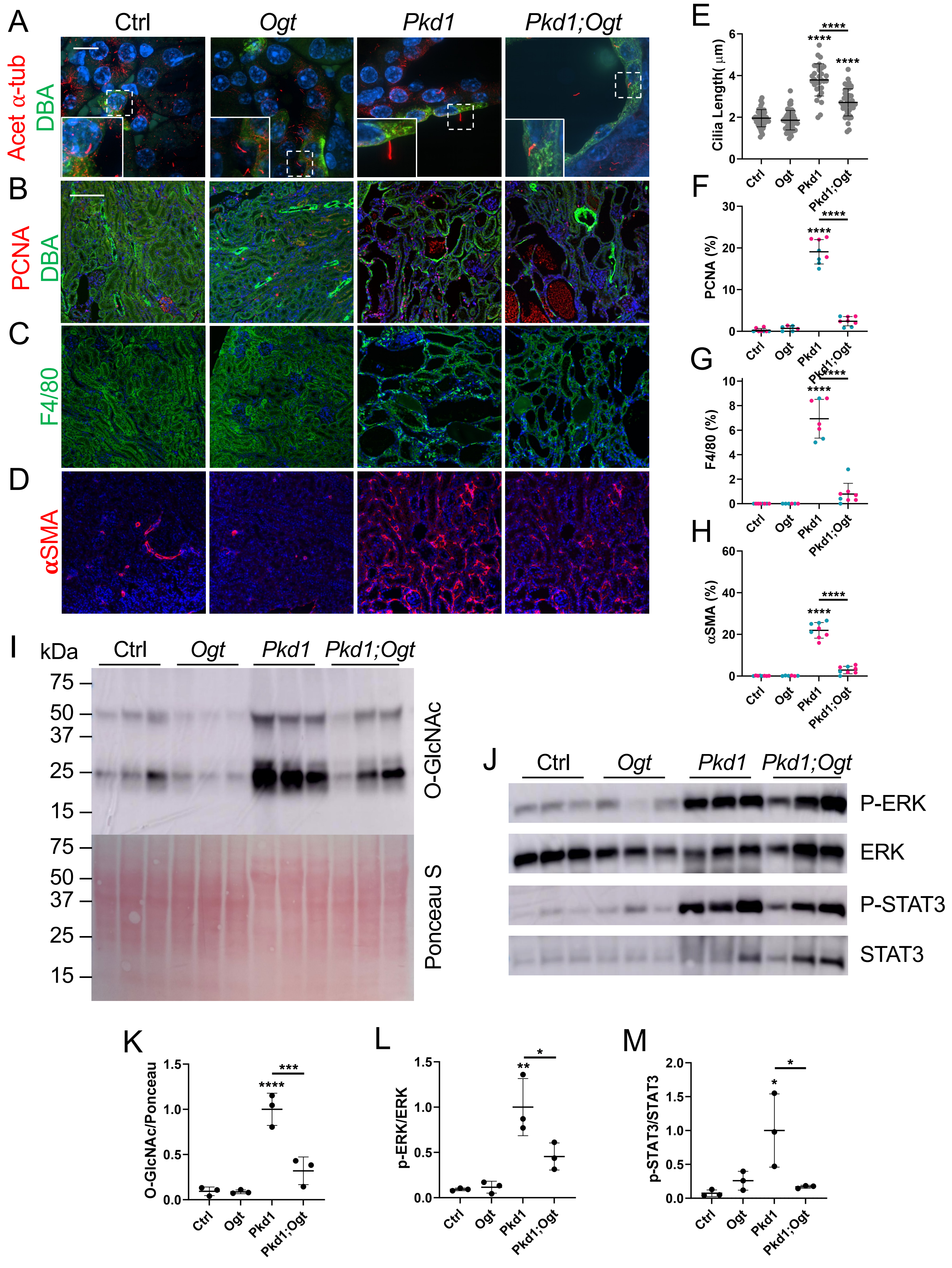
A
B
C
D
E
F
G
H
I
J
K
L
M
Ctrl
Ogt
Pkd1
Pkd1;Ogt
Acet α-tub
DBA
PCNA
DBA
F4/80
αSMA
Cilia Length( μm)
PCNA (%)
F4/80 (%)
αSMA (%)
****
kDa
75
50
37
25
15
O-GlcNAc
Ponceau S
P-ERK
ERK
P-STAT3
STAT3
O-GlcNAc/Ponceau
p-ERK/ERK
p-STAT3/STAT3
***
**
*

Figure S4

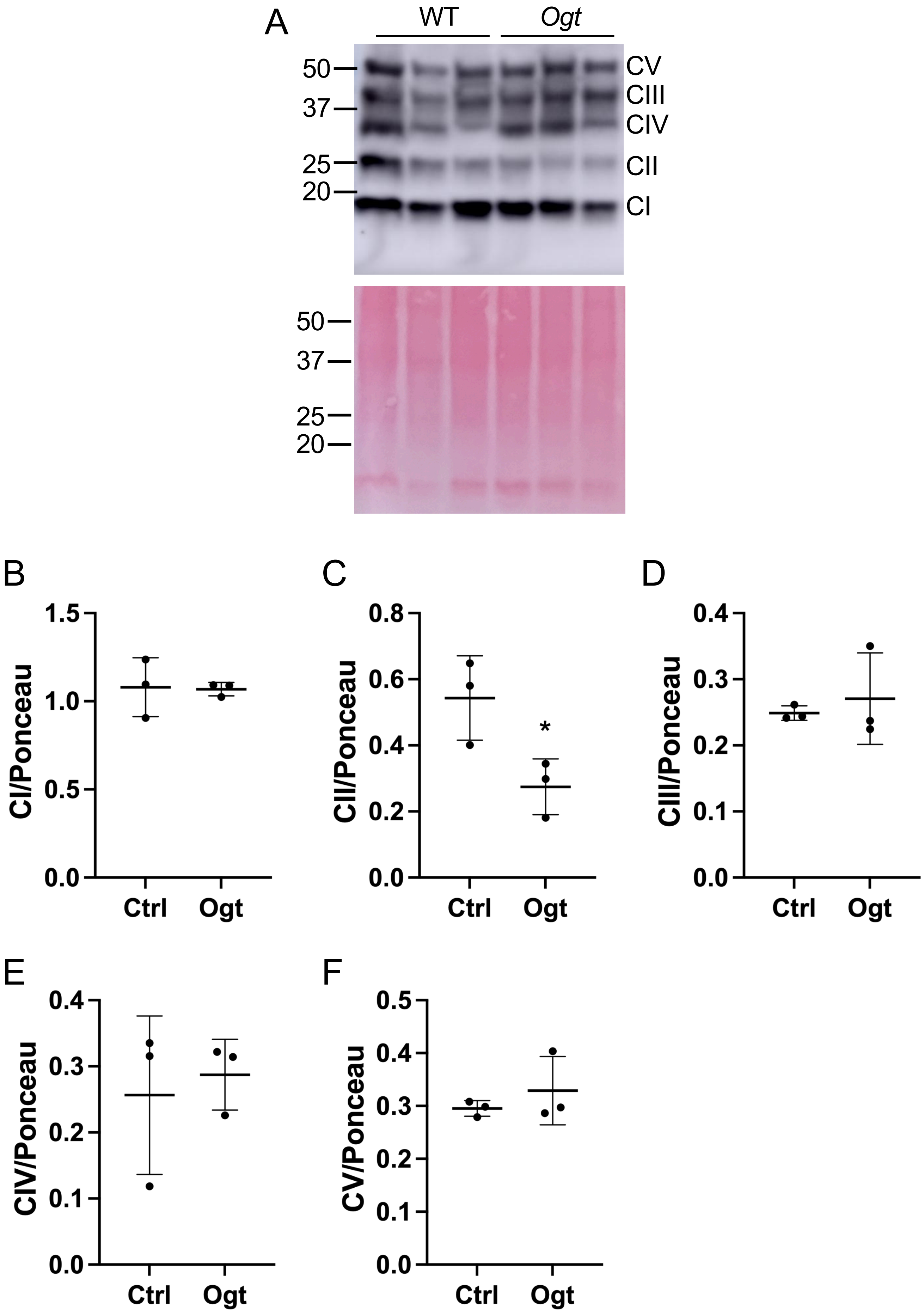

A
WT
Ogt
50
37
25
20
CV
CIII
CIV
CII
CI
B
CI/Ponceau
Ctrl
Ogt
C
CII/Ponceau
*
D
CIII/Ponceau
E
CIV/Ponceau
F
CV/Ponceau

Figure S5

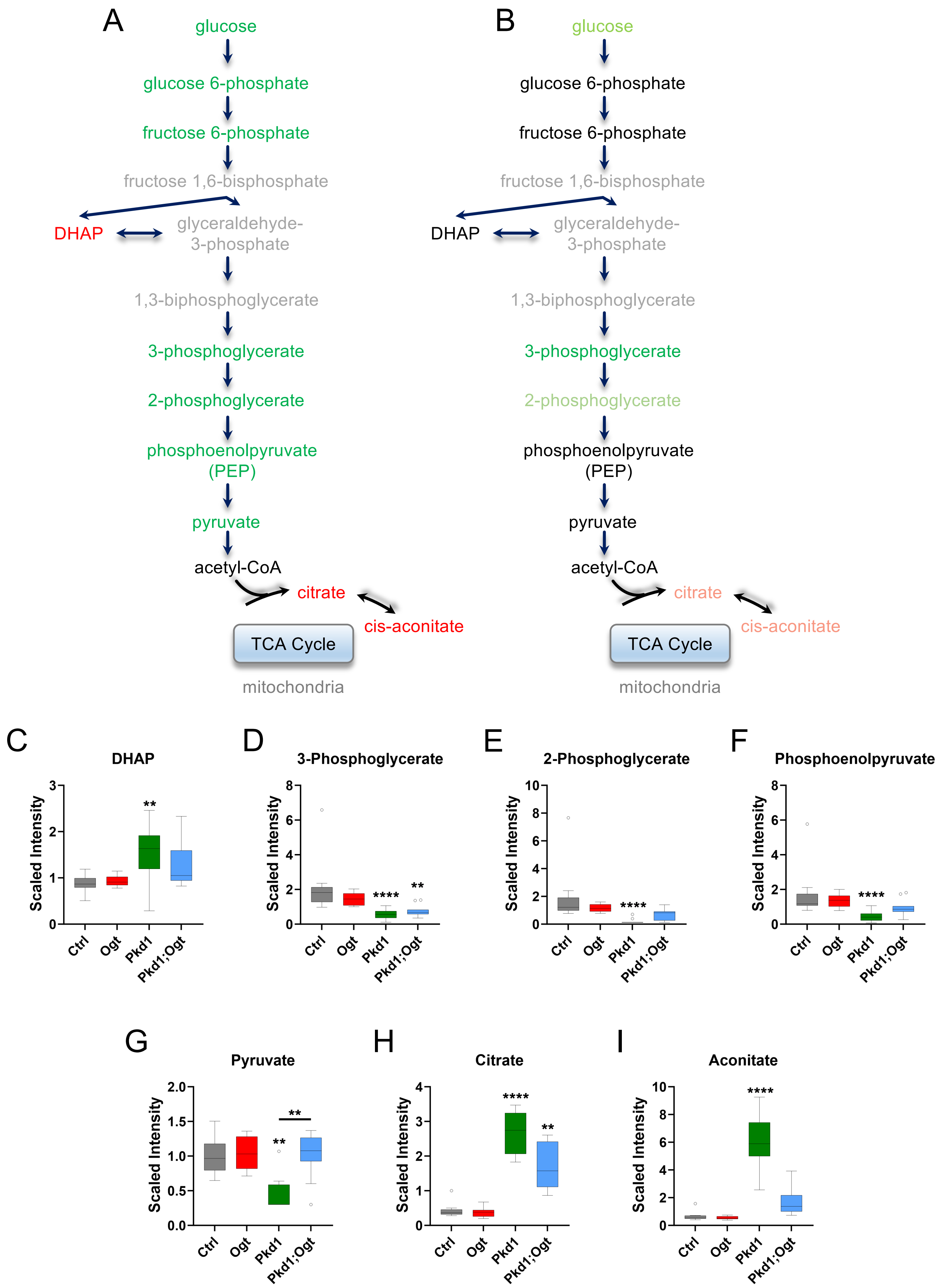
A
glucose
glucose 6-phosphate
fructose 6-phosphate
fructose 1,6-bisphosphate
DHAP
glyceraldehyde-3-phosphate
1,3-biphosphoglycerate
3-phosphoglycerate
2-phosphoglycerate
phosphoenolpyruvate (PEP)
pyruvate
acetyl-CoA
citrate
cis-aconitate
TCA Cycle
mitochondria
B
glucose
glucose 6-phosphate
fructose 6-phosphate
fructose 1,6-bisphosphate
DHAP
glyceraldehyde-3-phosphate
1,3-biphosphoglycerate
3-phosphoglycerate
2-phosphoglycerate
phosphoenolpyruvate (PEP)
pyruvate
acetyl-CoA
citrate
cis-aconitate
TCA Cycle
mitochondria
C
DHAP
D
3-Phosphoglycerate
E
2-Phosphoglycerate
F
Phosphoenolpyruvate
G
Pyruvate
H
Citrate
I
Aconitate
Scaled Intensity
Ctrl
Ogt
Pkd1
Pkd1;Ogt
**
****

### Figure S6

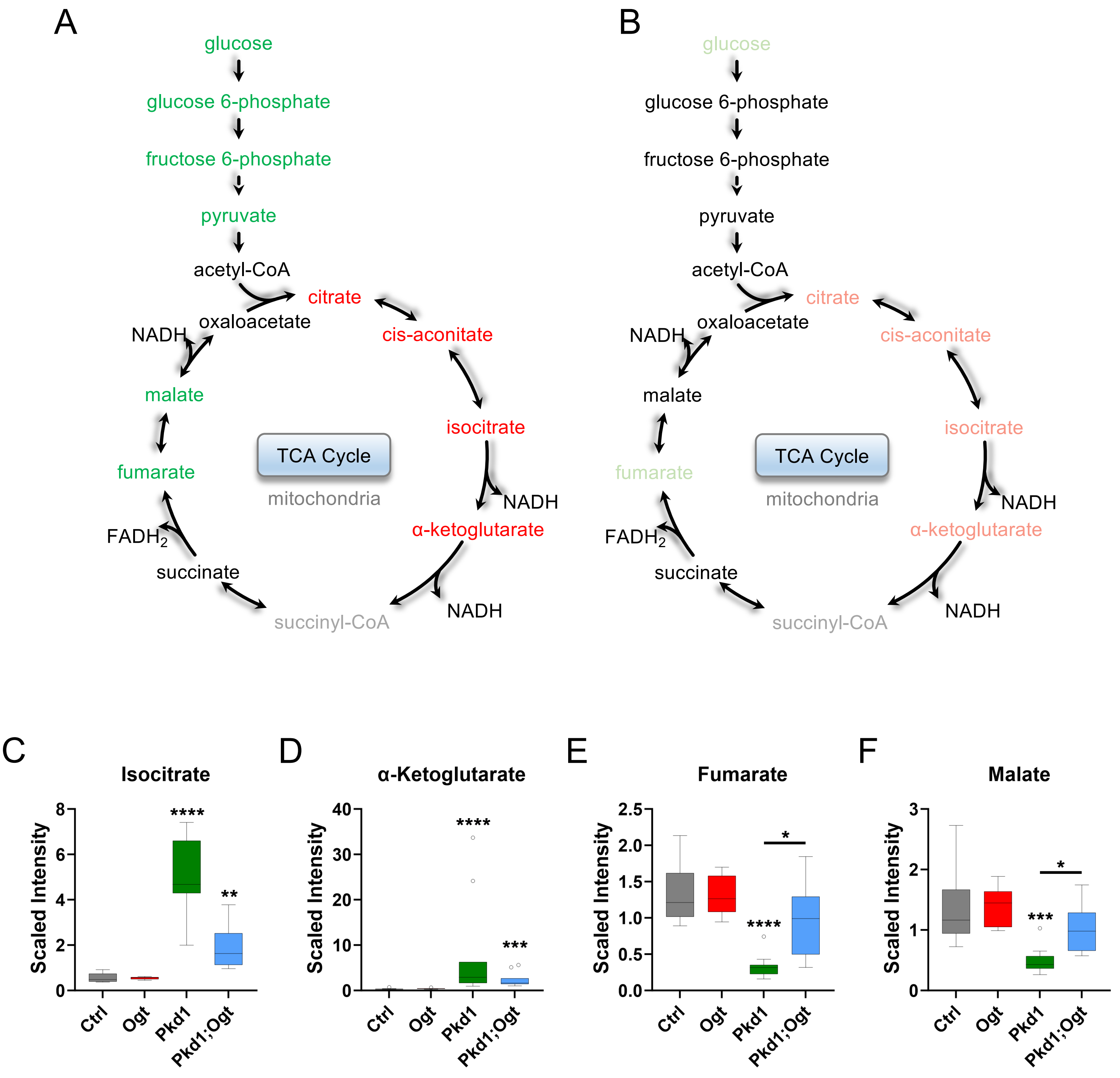

A
glucose
glucose 6-phosphate
fructose 6-phosphate
pyruvate
acetyl-CoA
citrate
oxaloacetate
cis-aconitate
NADH
malate
isocitrate
TCA Cycle
mitochondria
fumarate
NADH
α-ketoglutarate
FADH2
succinate
NADH
succinyl-CoA
B
glucose
glucose 6-phosphate
fructose 6-phosphate
pyruvate
acetyl-CoA
citrate
oxaloacetate
cis-aconitate
NADH
malate
isocitrate
TCA Cycle
mitochondria
fumarate
NADH
α-ketoglutarate
FADH2
succinate
NADH
succinyl-CoA
C
Isocitrate
Scaled Intensity
Ctrl
Ogt
Pkd1
Pkd1;Ogt
D
α-Ketoglutarate
Scaled Intensity
Ctrl
Ogt
Pkd1
Pkd1;Ogt
E
Fumarate
Scaled Intensity
Ctrl
Ogt
Pkd1
Pkd1;Ogt
F
Malate
Scaled Intensity
Ctrl
Ogt
Pkd1
Pkd1;Ogt

Figure S7

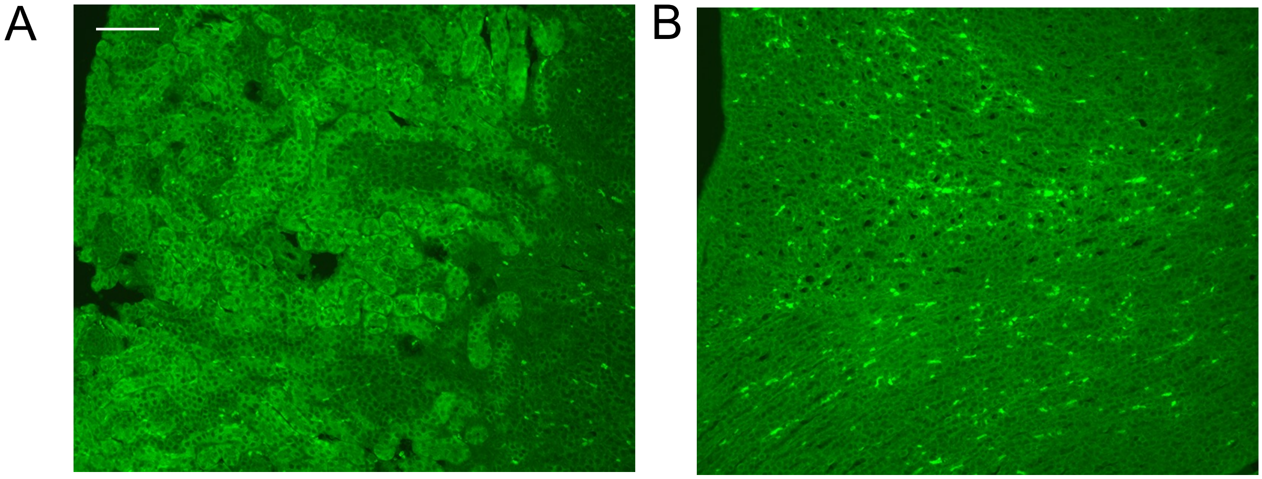
